# Epigenetic plasticity is associated with enhanced tolerance to low temperature stress in woodland strawberry

**DOI:** 10.64898/2026.04.24.719864

**Authors:** Relindis G. Njah, Stephen K. Randall, Jahn Davik, Wenche Johansen, Muath K. Alsheikh, Paul E. Grini, Robert C. Wilson

## Abstract

Low temperature stress causes significant damage to the strawberry plant. During cold stress, plants undergo morphological and physiological changes often regulated at the genetic and/or epigenetic levels. Some strawberry cultivars are more cold-hardy than others. Using the diploid woodland strawberry as a model, we analyzed the effects of cold acclimation on methylome and transcriptome dynamics in the crowns and leaves of three ecotypes with contrasting cold tolerance. ‘Alta’, which was the most cold-tolerant ecotype, exhibited the highest genetic and epigenetic plasticity in response to cold. CHH-context methylation dominated the differentially methylated regions (DMRs) with more hypomethylation in crowns and hypermethylation in leaves. CG methylation was enriched in gene bodies, while non-CG methylation was prevalent in upstream and downstream regions. Our study revealed that less than a quarter of differentially methylated genes (DMGs) showed changes in transcript accumulation levels. This finding indicates that universal cold response in *Fragaria vesca*, as reflected by gene expression, cannot be mechanistically attributed to DNA methylation. The majority of differentially expressed differentially methylated genes (DEDMGs) were ecotype- and tissue-specific. Enrichment analysis revealed that these genes were involved in pathways related to stress tolerance, such as carbohydrate metabolism, lipid metabolism, ATP hydrolysis, and cellular detoxification. Each ecotype responded to cold through mobilization of its own set of differentially expressed genes (DEGs), DMGs, and DEDMGs, and variation in expression and methylation patterns exhibited by ‘Alta,’ ‘FDP817’, and ‘NCGR1363’ suggest that cold signaling processes and survival depend on the tissue, ecotype, and geographical origin of the plants exposed to cold stress. Therefore, this study highlights the potential of both genetic markers and epialleles as molecular markers for the development of cold-tolerant octoploid strawberry cultivars that are better suited for propagation in Nordic climates.

## Introduction

Climate change has led to increasingly frequent and variable weather fluctuations, which pose major challenges for crop plants to remain healthy and productive. Frost-related low temperature stress is a major challenge for strawberry cultivation across the Northern hemisphere, affecting fruit quality, yield, harvest revenues, and the economic sustainability of production (Koehler et al. 2012; Davik et al. 2000). Phenotypical symptoms of cold stress in strawberry plants include tissue damage, necrosis, and reduced vigor (Nestby et al. 2000). This damage limits large-scale production, causing substantial economic losses for strawberry farmers in northern climates.

In response to cold stress, plants undergo physiological changes that can be regulated both by genetic and epigenetic mechanisms as part of their survival strategy. Genetic regulation under cold stress has been well characterized in model plants such as *A. thaliana*. Cold-regulated components of signaling pathways include membrane receptors, protein kinases, transcription factors, and other cold-responsive genes (Jeon and Kim 2013; Huang et al. 2012; Sun et al. 2019; Gusain et al. 2023). Crops such as strawberry, tomato, apple, wheat, tobacco, and rice exhibit cold-induced regulation of target genes following cold acclimation (Zhang et al. 2019; Zhao et al. 2018; Zhang et al. 2024, 2021; Sambe et al. 2015; Skinner 2009; Yang et al. 2023).

As temperatures approach freezing, plants activate various pathways that ultimately influence survival. Some of these pathways include the regulation of cold-induced genes at the transcriptional, post-transcriptional, translational, and post-translational levels (Gusain et al. 2023). Two major signaling pathways associated with cold stress are CBF-(C-repeat Binding Factor) dependent and CBF-independent. The ICE (Inducer for C-Repeat binding)-CBF-COR (COld-Regulated) signaling pathway is arguably the most studied pathway for protecting plants against low-temperature stress (D.-Z. Wang et al. 2017). ICE1 is a MYC-like transcription factor that binds DNA regulatory elements in the promoters of *CBF3/DREB1A* genes, thereby inducing *CBF* transcription. *CBF3/DREB1A*, an AP2/ERF-type transcription factor, subsequently regulates *COR* gene expression and contributes to cold tolerance (Qian et al. 2024).

Mutations in genes associated with CBF-dependent cold signalling alter plant responses to cold. Overexpression of *FvICE1* in woodland strawberry enhances cold tolerance and upregulates stress-responsive genes, whereas *fvice1* knock-out mutants exhibited increased cold sensitivity and reduced expression of these genes (Han et al. 2023). In *Arabidopsis*, cold-treated *ice1* exhibited reduced freezing tolerance and decreased *CBF3* levels, whereas overexpression of *ICE1* was associated with *CBF3* expression and improved freezing tolerance (Chinnusamy et al. 2003). In another study, a *cbf1 cbf2 cbf3* triple mutant of *A. thaliana* was highly sensitive to freezing after cold acclimation (Jia et al. 2016). *A. thaliana* plants lacking functional *OST1* (OPEN STOMATA 1) were hypersensitive to freezing stress (Ding et al. 2015). OST1 is a serine/threonine protein kinase whose activation by cold leads to *ICE1* phosphorylation and increased cold tolerance. Cold-activated *OST1* also phosphorylates AtANNEXIN1 (AtANN1), a plasma membrane-localized calcium transporter, thereby enhancing its Ca^2+^ transport activity. Loss-of-function *atann1* mutants exhibit reduced cold tolerance due to decreased cold-induced Ca^2+^ influx (Liu et al. 2021).

Genes and hormones involved in mediating cold stress response through the CBF-independent pathway include abscisic acid (ABA), *GIGANTEA* (*GI*), and *ESKIMO1* (*ESK1*) (Xin et al. 2007; Li et al. 2021; Cao et al. 2005). *ESKIMO1* is a negative regulator of cold stress through a CBF-independent pathway, and its mutation in *Arabidopsis* is associated with increased freezing tolerance (Xin et al. 2007). Loss-of-function mutations in *GI* resulted in reduced freezing tolerance (Cao et al. 2005).

The role of DNA methylation-based epigenetic regulation in cold response remains poorly understood. In plants, DNA methylation occurs in three sequence contexts: CG, CHG, and CHH, where H represents A, T, or C (Law and Jacobsen 2010). CG methylation within gene bodies has limited to no effect on gene expression, whereas CG methylation in 5’- and 3’-regulatory regions can inhibit transcription initiation by blocking the binding of RNA polymerase II and transcription factors (Zilberman et al. 2007; Zemach et al. 2010). Loss of non-CG methylation has been associated with loss of transposable element (TE) silencing, leading to defects in plant growth and development (Cheng et al. 2015; Hu et al. 2021). In plants, establishment of methylation in all sequence contexts is mainly carried out by DOMAINS REARRANGED METHYLTRANSFERASE 2 (DRM2) (Zhang et al. 2018; Law and Jacobsen 2010), while maintenance is mediated by METHYLTRANSFERASE 1 (MET1) in the CG context (Jones et al. 2001) and by CHROMOMETHYLASE 2 and 3 (CMT2 and CMT3) in the CHG context (Lindroth et al. 2001; Stroud et al. 2014). Asymmetric CHH methylation depends on continual *de novo* methylation mediated by DRM2 (Cao and Jacobsen 2002), primarily through the RNA-directed DNA methylation (RdDM) pathway. RdDM is a plant-specific process in which siRNAs generated from RNA polymerase IV (Pol IV) and PolV transcripts guide DRM2 *de novo* DNA methylation (Wang et al. 2023; Loffer et al. 2022; Zhang and Zhu 2025; Cao and Jacobsen 2002).

The impact of cold acclimation on DNA methylation has been reported in several plant species. For example, an overall decrease in DNA methylation levels was detected in cold-stressed *F. vesca* and sugar beet (Gutschker et al. 2022; López et al. 2022), whereas an increase in global DNA methylation was identified in *F x ananassa* and *A. thaliana* in response to cold (Zhang et al. 2012; Rahman et al. 2024). In apple, cold stress resulted in a decrease in total DNA methylation, while no significant reduction was detected in plants exposed to milder chilling temperatures (Kumar et al. 2016). In rice, cold-induced DNA methylation changes depended on growth stage or organ or tissue type, and ecotype (Pan et al. 2011). Together, these studies reveal inconsistent global DNA methylation responses to cold, suggesting that the underlying regulatory mechanisms may act at single genes rather than at the whole-genome level.

Stress-induced changes in DNA methylation alter gene expression and influence phenotype. In *A. thaliana,* hypermethylation of *ICE1* is associated with reduced *CBF* expression and decreased cold tolerance (Xie et al. 2019). In roses, cold-induced DNA hypermethylation in the promoters of scent-related genes and of the *AGAMOUS* homolog, *RhAG*, correlates with their downregulation, accompanied by reduced fragrance and an increase in petal number (Ma et al. 2015; Xie et al. 2023). In peach, loss of flavour has been associated with promoter hypermethylation of flavor-related genes resulting in their reduced expression (Duan et al. 2023). However, there is still limited evidence that DNA methylation at individual genes regulates cold-responsive gene expression in strawberry.

Strawberry crowns and leaves play important roles in surviving low-temperature stress. Leaves, as primary photosynthetic organs, are directly exposed to environmental stress, and thus are critical sites of signal perception and transduction (Zhang et al. 2019). The crown serves as a carbohydrate storage organ, providing energy for flowering and fruit development. Although highly sensitive to cold stress, crown viability is essential for winter survival, regeneration of spring leaves, and yield potential (Macías-Rodríguez et al. 2002; Shokaeva 2008).

In this study, we have investigated the impact of cold treatment on DNA methylation dynamics and transcript accumulation in crowns and leaves of three *Fragaria vesca* (*F. vesca*) ecotypes with large variation in cold susceptibility. We asked whether low-temperature stress induces changes in DNA methylation and gene expression, and how cold-induced methylation patterns correlate with gene expression within and between ecotypes. Our results reveal higher basal methylation levels and a more pronounced combined response of DNA methylation and gene expression in the most cold-tolerant ecotype, which may collectively or individually contribute to enhanced freezing tolerance. Furthermore, we identified genes exhibiting differential DNA methylation in response to cold as a more precise alternative that were regulated similarly across all ecotypes, suggesting an important canonical role of DNA methylation in cold adaptability in *F. vesca*.

## Material and methods

### Plant material and treatment conditions

This study was conducted using the freezing-tolerant Norwegian ecotype ‘Alta’ from northern Norway, the moderately cold-tolerant ecotype ‘NCGR1363’ from Bolivia, and the cold-susceptible ecotype ‘FDP817’ from California (Table S1). Each ecotype was propagated through runners to produce a set of test plants. Runners were rooted and grown in 10 cm plastic pots containing a peat-based compost (90 % peat, 10 % clay), with the addition of 1:5 (v/v) of granulated perlite raised in a greenhouse for five weeks at 20 ± 2 °C under an 18-hour photoperiod. The plants were watered twice a week with a balanced nutrient solution containing 7.8 mmol N, 1 mmol P, and 4.6 mmol K per litre.

After five weeks in a common garden, half the plants remained in the greenhouse (untreated), and half were kept in a cold room at 2 °C under a 12-hour photoperiod (cold-treated) for 42 days. Plants were watered with cold water as needed. After 42 days, three biological replicates were harvested from cold-treated (CT) and untreated (UT) crown and leaf tissues of each ecotype, immediately frozen in liquid nitrogen and stored at −80 °C prior to DNA or RNA extraction. Untreated plants were harvested in the greenhouse, and cold-treated plants were harvested in the cold room. Each biological replicate comprised whole plant leaves and entire crown tissue from five plants of uniform size and similar developmental stage. Although collected from different mother plants, the biological replicates are genetically identical because all mother plants were propagated from a single parent.

### Library construction and bisulfite sequencing

Genomic DNA was isolated from three biological replicates of both crown and leaf tissue from each ecotype using the DNAeasy Plant Kit (Qiagen). To meet the minimum DNA input required for bisulfite library construction, equimolar amounts of three biological samples per treatment were pooled to a final concentration of 5 μg. Samples with a 260/280 ratio > 1.8 were normalized to 2.5 ng/μl using a Qubit fluorometer (Thermo Scientific). Whole genome bisulfite sequencing library preparation was performed by the Splinted Ligation adapter Tagging method (Raine et al. 2016). Genomic DNA was sheared to 300 bp (Covaris, M220 Focused-ultrasonicator and Holder XTU) and fragments treated with sodium bisulfite (Zymo EZ Methylation Gold kit, Zymo Research), end-repaired, 5’-phosphorylated by polynucleotide kinase (Thermo Scientific), 3’-adenylated with Taq DNA polymerase, and ligated to splinted adapters. Libraries were amplified using the KAPA HiFi Uracil+ PCR 2x Master Mix (KAPA Biosystems, USA) and Illumina PCR primers. Twelve libraries (two tissue types × three parental ecotypes × two timepoints) of 150 bp paired-end reads were sequenced on an Illumina HiSeq 3000 instrument (Norwegian Sequencing Centre, University of Oslo). PhiX genomic DNA (20 %) was added to each lane of the flow cell to increase sequence diversity, quality, and calibration control.

### RNA Sequencing

Crown and leaf tissue from ecotypes ‘Alta,’ ‘FDP817,’ and ‘NCGR1363,’ either exposed to low temperature stress (2 °C for 42 days; cold-treated) or maintained as controls (20 ± 2 °C; untreated), were pulverized in liquid nitrogen, and RNA was extracted using the Spectrum Plant Total RNA Kit (Sigma). Thirty-six libraries (two tissue types × three parental ecotypes × two timepoints × three biological replicates) were prepared from RNA samples with a RIN (RNA Integrity Number) >8 using the strand-specific TruSeq™ RNA-seq library preparation kit (Illumina), and 150 bp paired-end reads were generated over three lanes of an Illumina HiSeq 4000 instrument (Norwegian Sequencing Centre, University of Oslo).

### Bioinformatics and statistics

Quality control of reads before and after trimming was performed using FastQC (Version 0.11.7, www.bioinformatics.babraham.ac.uk/projects/fastqc/). Low-quality reads were filtered and adapter sequences trimmed using Trim Galore (version 0.4.4_dev, www.bioinformatics.babraham.ac.uk/projects/trim_galore/). Bismark (Krueger and Andrews 2011) was used to map trimmed reads to the *Fragaria vesca* (of the Hawaii 4 accession) assembly version 4.0 (FvH4.0.a2; (Edger et al. 2018; Li et al. 2019) reference genome using Bowtie 2 algorithm, remove duplicated reads (Bismark deduplication step) and generate -CX report files (Bismark methylation extractor step) used as the basis for calling differentially methylated regions (DMRs).

DMRs were identified by comparing methylation levels of untreated and cold-treated samples of each ecotype, using the DMRcaller pipeline (Catoni et al. 2018). DMRs were called by comparing the number of methylated cytosines in UT and CT samples using the ‘bins’ method, with *binSize* 100 bp for CG and CHG contexts and 50 bp for the CHH context. A DMR was required to contain a minimum of four cytosines, each covered by at least four reads, and each bin was required to have a methylation proportion (methylated cytosines / all cytosines) difference of at least 0.2 and a Benjamini-Hochberg adjusted p-value threshold of 0.01 (Score test). The parameter *minGap* was set at 0 bp to prevent merging of DMR bins, and *minSize* was set to 0. Variation in Global DMR patterns between ecotypes and tissues was visualized using the Circos software package (Krzywinski et al. 2009). DMRs were annotated using the Splicejam package in R (github.com/jmw86069/splicejam). Bar plots and scatterplots were generated using easyGgplot2 and ggplots, respectively. Venn diagrams were generated using a web-based tool (bioinformatics.psb.ugent.be/webtools/Venn/).

MethPipe (Song et al. 2013) was used to generate data showing average global methylation patterns in genic (GE) and TE regions for untreated and cold-treated samples. Weighted DNA methylation averages (Schultz et al. 2012) were calculated across genomic regions using the ‘roimethstat’ module of Methpipe, and the output was used to generate metaplot data showing methylation proportions. Specifically, genic and TE elements (upstream, body, downstream) were each divided into 20 bins, proportionally for gene and TE bodies that inherently vary in length, and the average DNA methylation level for each such bin across all respective genomic regions was calculated (github.com/Njah/Njah-et-al.-Manuscript-2026) before generating plots in R using the ggplot2 package. The upstream and downstream regions in GE and TE refer to 1 kb upstream of transcription start sites (TSS) and 1 kb downstream of transcription end sites (TES), respectively. Circos plots describing absolute methylation at the chromosome level were generated using Shinycircos (Yu et al. 2018). Linear representation of absolute methylation was generated using DMRcaller.

Transcriptomics analysis was performed using the OmicsBox (version 2.1.2) platform. Trimmed reads were mapped to the FvH4.0.a2 transcriptome reference (https://www.rosaceae.org/) using RSEM (version 1.3.2; (Li and Dewey 2011) and Bowtie2 (Langmead and Salzberg 2012). Differentially expressed genes were identified using the OmicsBox edgeR Bioconductor package (Robinson et al. 2010)version 3.28.0; (Robinson et al. 2010) with the following parameters: FDR cut off 0.05; generalized linear model (GLM) quasi-likelihood F-test; CPM cutoff 0.5 in a minimum of 2 of 3 biological replicates; and sample normalization based on TMM. The threshold for calling a DEG was set to a fold change (FC) > 2 or FC < −2 with p.adjust < 0.05 for upregulation or downregulation, respectively. For quality control, sample homogeneity was assessed by principal component analysis (PCA) using the DESeq2 package (Love et al. 2014). The probability of overlap between DMGs and DEGs was assessed using an online tool running the hypergeometric test (nemates.org/MA/progs/overlap_stats.html). Gene ontology (GO) term enrichment analysis was performed using ClueGO, a Cytoscape plug-in (Shannon et al. 2003; Bindea et al. 2009) with an FDR-adjusted p-value < 0.01.

## Results

### Cold stress altered DNA methylation in CHG and CHH contexts

Plant runners from ‘Alta,’ ‘NCGR1363’ and ‘FDP817’ were rooted and grown in a greenhouse for five weeks at 20 ± 2 °C and an 18-hour photoperiod. After five weeks, half of the population was acclimated at 2 °C for 42 days, and the other half (controls) remained in the greenhouse during the cold acclimation period. To study the DNA methylation patterns, DNA from crowns and leaves of untreated and cold-treated plants was bisulfite-treated, sequenced, and mapped to the *Fragaria vesca* reference genome. Mapping efficiency ranged from 52 % to 71 % for bisulfite-converted DNA (Table S2). To investigate the impact of cold treatment on DNA methylation in ‘Alta,’ ‘FDP817’ and ‘NCGR1363,’ we compared global cytosine methylation patterns in crowns and leaves of untreated and cold-treated samples. For brevity, we most often emphasize results in crowns, which are the central tissue important for overwinter survival in *Fragaria* (Shokaeva 2008).

A global methylation response to cold stress was observed in all three ecotypes. Similar methylation landscapes between ecotypes and distinct chromosome-specific methylation patterns were also identified (Fig. 1A, Fig. S1). However, characteristic ecotype-specific methylation patterns were identified, primarily in ‘Alta’ (Fig. S2 and Fig. S3). By direct comparison of methylation levels among ecotypes, ‘Alta’ displayed higher methylation levels than corresponding chromosomal regions in ‘FDP817’ and ‘NCGR1363’ (Fig. S4, S5, S6, and S7).

**Figure 1.**
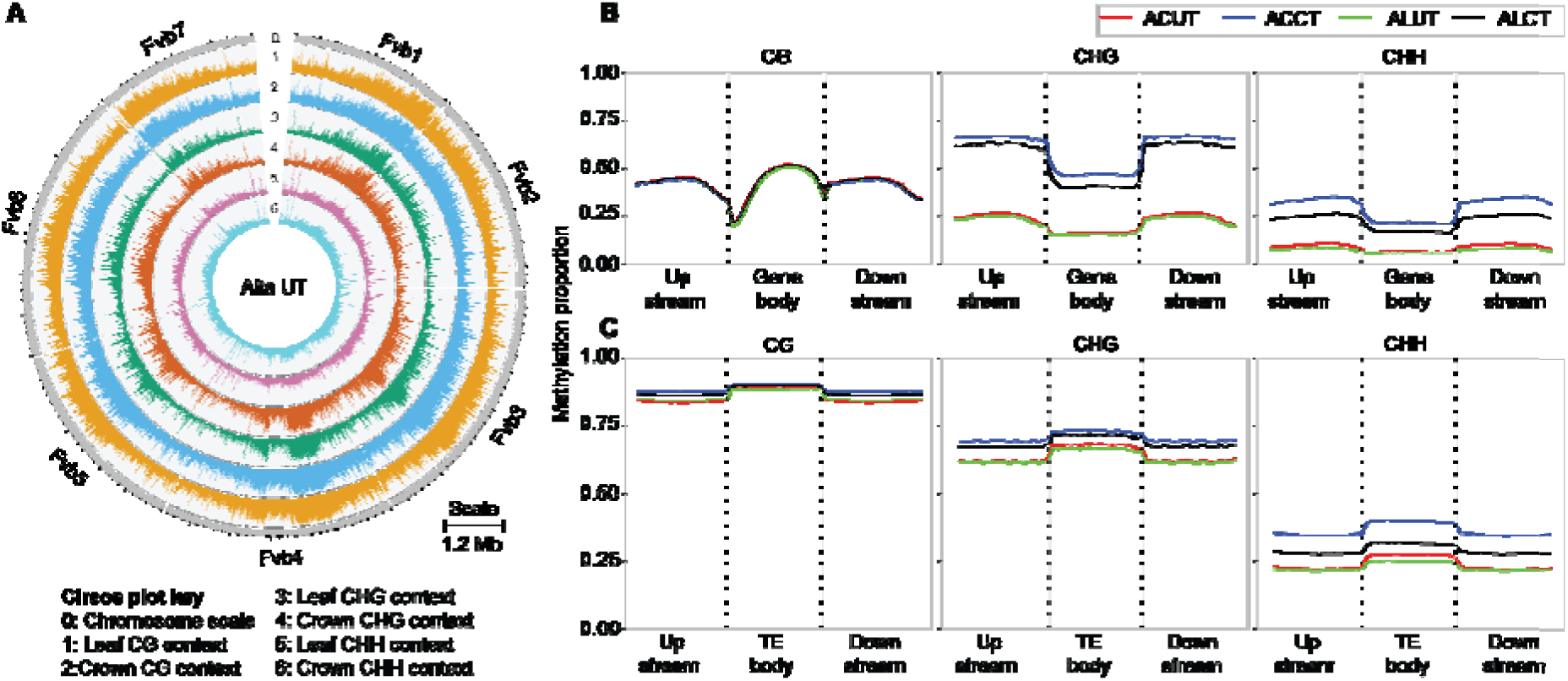
Global DNA methylation landscape in crowns and leaves of ‘Alta.’ (A) Circos plot shows methylation patterns in CG, CHG, and CHH contexts of all seven chromosomes of ‘Alta’ crowns and leaves propagated at normal (UT; 20 ± 2 °C) temperature conditions. Lanes 1-6 show methylation levels (methylated reads/total reads) for each tissue and context. Methylation levels are shown for all methylation contexts along all seven F. vesca chromosomes, using a 50 kb window. (B) DNA methylation patterns in CG, CHG, and CHH sites of genic regions in ‘Alta’ crowns (AC) and leaves (AL), propagated at normal (20 ± 2 °C) and low (CT; 2 °C) temperatures for 42 days. (C) DNA methylation patterns in CG, CHG, and CHH sites of TE regions in ‘Alta’ crowns (AC) and leaves (AL), propagated at normal (20 ± 2 °C) and low (CT; 2 °C) temperatures for 42 days. Plots show cold-induced methylation changes in CHG and CHH contexts in both genic- and TE regions. TE regions have higher basal methylation levels (UT) and exhibit a lower degree of methylation changes after cold acclimation compared with genic regions. Upstream and downstream regions were defined as 1 kb in size. Colored lines show different combinations of tissue and treatment: ACUT = ‘Alta’ crown untreated; ACCT= ‘Alta’ crown cold-treated; ALUT = ‘Alta’ leaf untreated; ALCT= ‘Alta’ leaf cold-treated.

Global analysis of methylation patterns in genic regions across all three contexts showed higher average levels in CG and CHG than in CHH. The distribution of DNA methylation peaked around upstream and downstream regions in CHG and CHH contexts and exhibited higher methylation levels in flanking regions than in gene bodies, where CG methylation was higher but dropped sharply towards their 5’ ends. Cold treatment induced global DNA methylation changes in crowns to a greater degree than in leaves at CHG and CHH sites, with the lowest methylation levels observed in leaves (Figure 1B; Fig. S8A, C). Analysis of TE methylation also showed high basal methylation levels (i.e., in untreated plants) and lower cold-induced methylation compared with genic regions (Figure 1C; Fig. S8B, D).

### Numbers of differentially methylated regions (DMRs) in genic regions varied among ecotypes and were pervasive in CHH sites

Next, we identified differentially methylated regions (DMRs) between untreated and cold-treated plants. In crown tissue, the numbers of identified DMRs varied between ecotypes (ranges of 281-2149 in CG, 311-2707 in CHG, and 9709-18337 in CHH), and similar variation was also found for leaves (643-2413 in CG, 680-1912 in CHG, and 7634-18999 in CHH; Table S3, SData 1). ‘Alta’ had 88 % and 15 % more DMRs than ‘FDP817’ and ‘NCGR1363,’ respectively. Crowns of ‘Alta’ and ‘FDP817’ exhibited higher numbers of DMRs than their corresponding counts in leaves, whereas NCGR1363 showed the opposite pattern (Figure 2A; Fig. S9A; Table S3). Across ecotypes, methylation was most prevalent in the CHH context.

**Figure 2.**
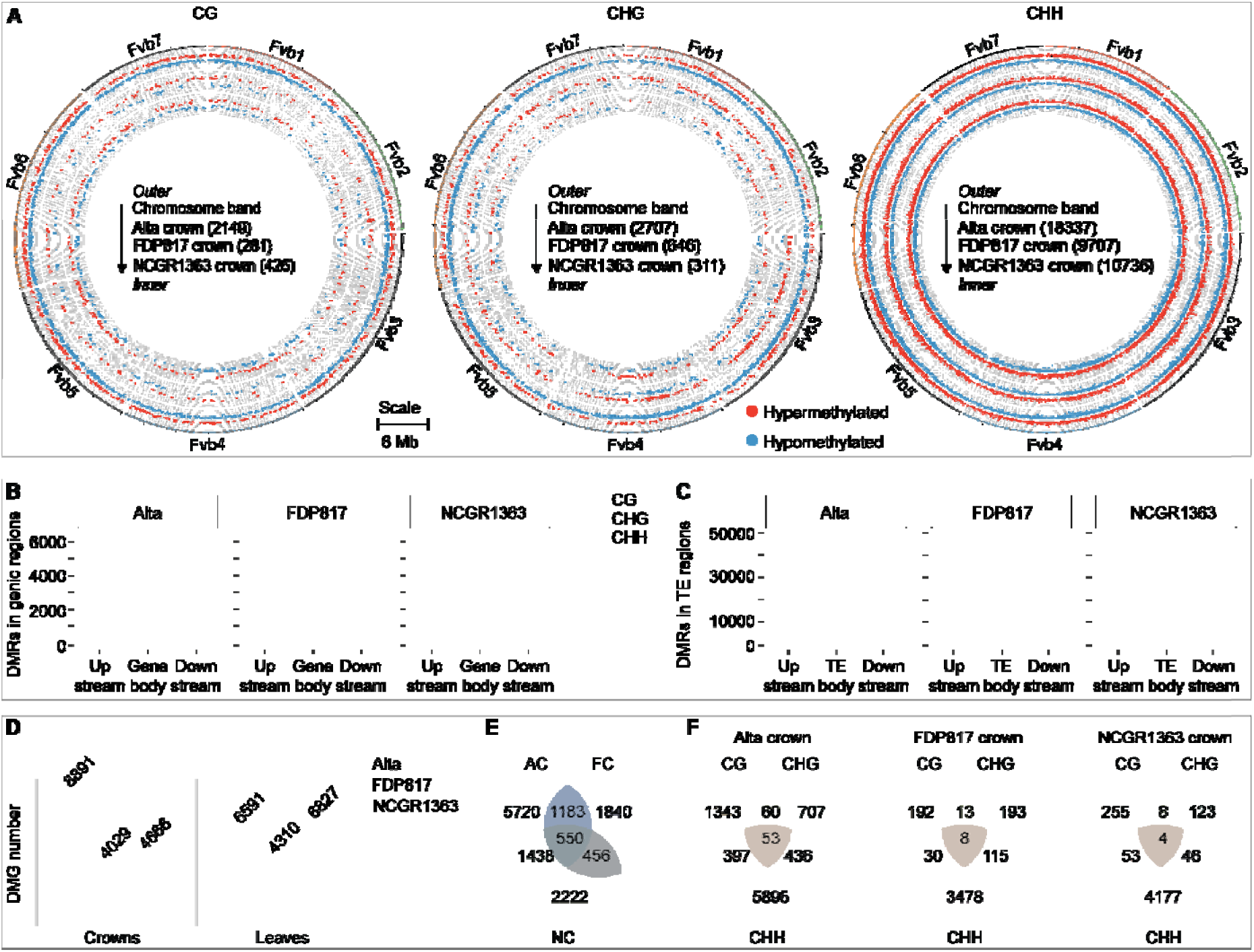
Differential methylation in genic elements and contexts of ‘Alta,’ ‘FDP817’ and ‘NCGR1363’ ecotypes. (A) Distribution of differentially methylated regions in CG, CHG, and CHH contexts in all seven chromosomes of ecotype crowns. ‘Alta’ has the highest number of DMRs in all methylation contexts. Numbers of DMRs in each tissue are indicated in parentheses. Red and blue dots denote hypermethylated and hypomethylated regions, respectively. (B) Numbers of DMRs overlapping with CG, CHG, and CHH sites in genic regions in crowns of all three ecotypes. (C) Numbers of DMRs overlapping with CG, CHG, and CHH sites in TE regions in crowns of all three ecotypes. ‘Alta’ displayed the highest number of DMRs across genomic regions. (D) Total numbers of DMGs in crowns and leaves of all three ecotypes. (E) Numbers of DMGs that were either unique to one ecotype or shared with others. Over half of DMGs in ‘Alta’ were unique, while ‘FDP817’ and ‘NCGR1363’ shared a majority of their DMGs. (F) Numbers of genes with DMRs in either one or multiple methylation context(s). A majority of genes were methylated exclusively in one context.

Focusing on genic regions, DMRs were evenly distributed across all three genic regions in all three ecotypes (Figure 2B; Fig. S9B; SData 2). The absolute numbers of DMRs overlapping genic regions were much higher in Alta crowns, with a greater proportion in the CHG and CHH contexts compared with the two other ecotypes. The number of DMRs in TE regions was close to ten times higher than in genic regions (Figure 2C; Fig. S9C; SData 3) and was most pervasive in the CHH context. CG methylation was more prevalent in gene bodies and in regions flanking TE bodies, whereas the opposite methylation pattern was observed for CHG and CHH contexts (Table S4 and Table S5).

### More genes were differentially methylated in ‘Alta’ and ‘NCGR1363’ than in ‘FDP817’

Genes whose upstream, downstream, and/or body regions overlapped DMRs were defined as differentially methylated genes (DMGs). At 8891, ‘Alta’ displayed roughly twice as many DMGs in crowns as each of the other two ecotypes (Figure 2D). In leaves, DMG numbers were more even across ecotypes, although ‘FDP817’ had lower numbers than the other two.

We investigated DMGs shared between ecotypes. Most DMGs in crowns of ‘Alta’ and in leaves of both ‘Alta’ and ‘NCGR1363’ were ecotype specific; ‘Alta’ and ‘NCGR1363’ also shared more DMGs than either did with ‘FDP817’ (Figure 2E; Fig. S10A; SData 4). Regarding methylation contexts, the overall majority of DMGs exhibited CHH methylation, with a subset also showing additional methylation at CG and CHG sites (Figure 2F; Fig. S10B). Notably, ‘Alta’ had over five times the number of DMGs exhibiting CG methylation compared with the two other ecotypes. Enrichment analysis revealed that DMGs were significantly (p.adjust < 0.01) enriched in nearly 140 GO terms (SData 5; Fig. S11). Although some terms were shared across contexts, tissues, and genic elements in selected ecotypes, these were not further analyzed.

### Differentially methylated genes are associated with gene regulation

To explore the association between DNA methylation and gene expression, differential gene expression analysis was performed (Table S6). RNA was extracted from crowns and leaves of untreated and cold-treated plants from the three ecotypes and paired-end RNA-seq reads were mapped to the second transcript annotation version of *Fragaria vesca* reference genome assembly version 4 (FvH4.0.a2) with mapping efficiencies ranging from 68 % to 82 % (Table S6). Principal component analysis (PCA) of normalized read counts showed high homogeneity among replicate samples within each treatment group (Figure 3A; Fig. S12A). Substantial differential gene expression between treated and untreated samples was observed in all ecotypes (Figure 3B; Fig. S12B; SData 6) and GO-term enrichment was detected among upregulated and downregulated DEGs in ecotypes (Figure 3C; SData 7).

**Figure 3.**
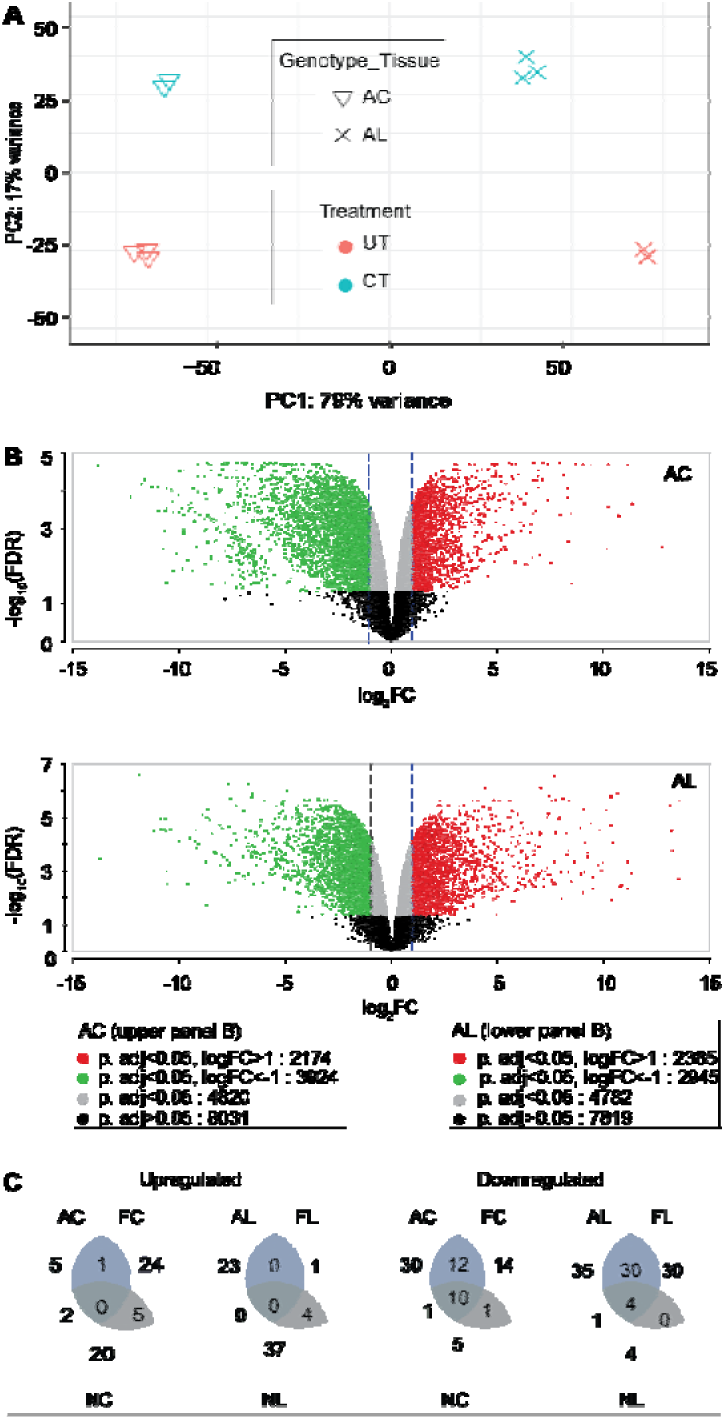
Characterization of transcriptional changes during cold acclimation. (A) Clustering of replicates into treatment groups for crowns and leaves of ‘Alta’ propagated at control (20 ± 2 °C) and cold acclimation (2 °C) temperatures for 42 days. Analysis revealed homogeneity within replicates of the same treatment group for all replicates used in the downstream analysis. (B) Volcano plots of expressed (-log_10_FDR) genes versus their degree of fold change (log_2_FC). Significantly differentially expressed genes (p.adjust < 0.05) are marked in green, red, and grey, with red and green denoting genes with log_2_FC values greater than 1 (upregulated) and less than −1 (downregulated), respectively. Black dots indicate non-significant (p.adjust ≥ 0.05) DEGs, while grey dots display significant DEGs. Points within dashed lines define −1 ≤ log_2_FC ≤ 1. (C) Numbers of enriched GO terms in upregulated and downregulated DEGs in crowns and leaves of ‘Alta,’ ‘FDP817’, and ‘NCGR1363.’ AC= ‘Alta’ crown, AL= ‘Alta’ leaf, FC= ‘FDP817’ crown, FL= ‘FDP817’ leaf, NC= ‘NCGR1363’ crown, NL= ‘NCGR1363’ leaf, UT= untreated, CT= cold-treated.

Among the identified DMGs, 8 % to 17 % were significantly differentially expressed (log_2_FC > 1 or log_2_FC < −1; p.adjust < 0.05). These were classified as Differentially Expressed Differentially Methylated Genes (DEDMGs; Figure 4A; Fig. S13A; Fig. S14A; Table S7A; SData 8), with ‘Alta’ crowns having the highest number. Although most DMRs in DMGs were specific to a genomic region (Figure 4B; Fig. S13B; Fig. S14B; Table S7B) or methylation context (Figure 4C; Fig. S13C; Fig. S14C), a substantial share of DEDMRs were also found to have DMRs in multiple genic regions or across methylation contexts. Although the observed DMG–DEG overlap deviates from random expectation by only about 10 %, the large sizes of the two datasets and the finite number of genes in the *F. vesca* genome make this deviation highly significant; therefore, the overlap likely reflects a real biological association (Table S7C).

**Figure 4.**
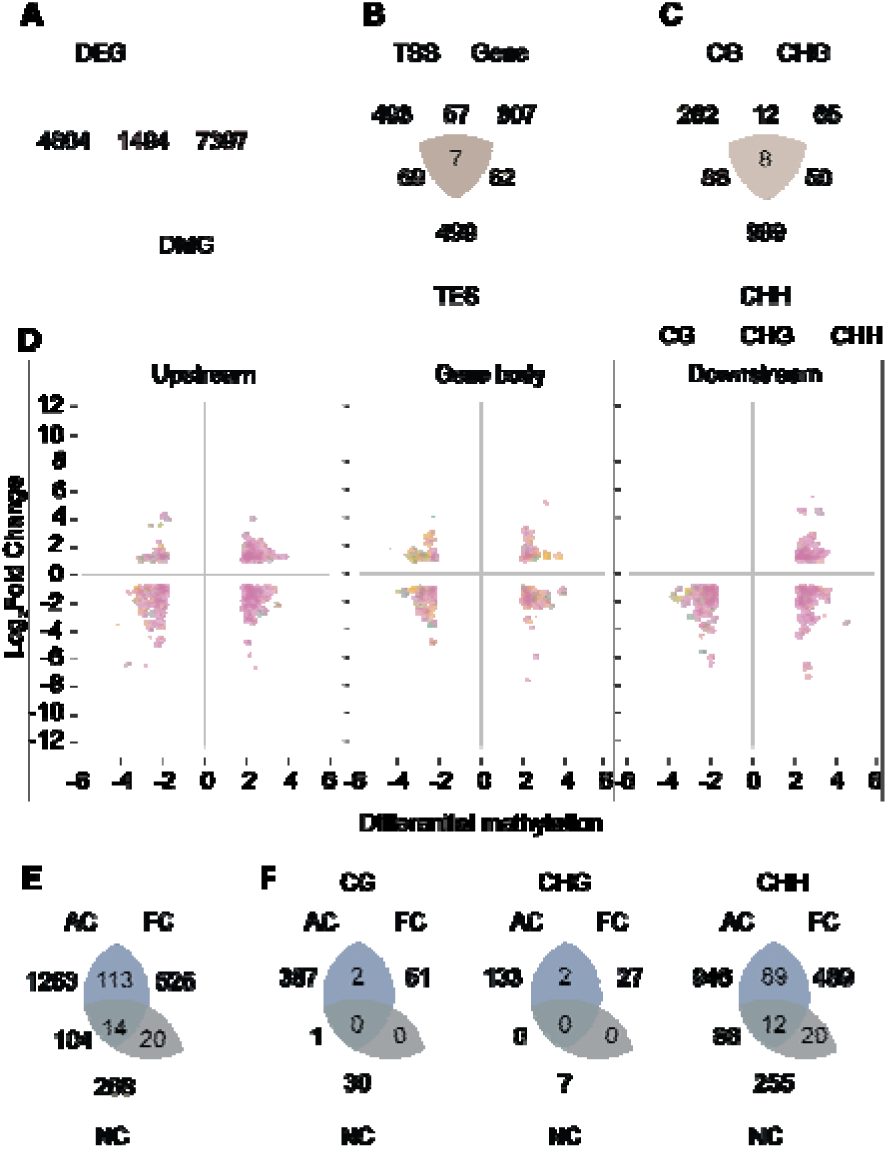
Differentially expressed differentially methylated genes in crowns. (A) Overlap between DEGs and DMGs in ‘Alta.’ (B) Numbers of DEDMGs harboring DMRs in either one or multiple genic regions. (C) Numbers of DEDMGs with DMRs in one or multiple methylation contexts. A majority of DEDMGs were methylated in one genic region or methylation context. (D) Differentially expressed genes associated with DMRs (DEDMRs), showing relationships between transcript accumulation (log_2_FC) and DNA methylation levels (DNA methylation proportion difference between cold-treated and untreated samples) in CG, CHG, and CHH contexts of gene bodies and regions 1 kb upstream of TSS and 1 kb downstream of TES in ‘Alta.’ (E) Numbers of shared and unique DEDMGs in ecotypes regardless of methylation context and region. (F) Numbers of shared and unique DEDMGs in CG, CHG, and CHH contexts among ecotypes. A majority of DEDMGs were ecotype-specific. AC= ‘Alta’ crown, FC= ‘FDP817’ crown, NC= ‘NCGR1363’ crown.

Integrative analysis of gene expression and methylation in genic regions of DEDMGs classified them into four groups: hypermethylated-upregulated, hypermethylated-downregulated, hypomethylated-upregulated, and hypomethylated-downregulated genes (Figure 4D; Fig. S13D-E; Fig. S14D-F; SData 9). ‘Alta’ had the greatest number of DMRs in all genic regions of DEDMGs. Most of these were downregulated and hypermethylated in the CHH context (SData 9).

### A majority of DEDMGs were ecotype-specific with prevalent CHH methylation

Depiction of crown DEDMGs across the three ecotypes demonstrated that most DEDMGs were ecotype-specific (Figure 4E). Over 65 % of DEDMGs were ecotype-specific across tissues, irrespective of methylation context or genomic region (Figure 4E, Fig. S16A, SData 10). In terms of methylation context, DEDMGs in CG or CHG contexts were predominantly ecotype-specific, whereas up to one third of CHH-methylated DEDMGs were shared among ecotypes (Figure 4F; Fig. S16B).

Enrichment analyses of DEDMGs unique to tissues of ecotypes identified 39 GO terms, all of which were tissue-specific (Fig. S15; SData 11D). Enriched GO terms included ‘transcription regulation,’ ‘oxidoreduction activity,’ and ‘carbohydrate metabolism’ in ‘Alta’ crowns. Genes with family members previously associated with cold stress acclimation were identified within these GO terms. For ‘DNA-binding transcription factor activity,’ genes were found from the ethylene□responsive transcription factor family, the WRKY family of transcription factors, one gene encoding a *MADS*-box transcription factor family member, a *DEHYDRATION-RESPONSIVE ELEMENT-BINDING PROTEIN 1E* (*DREB1E*) gene, all exhibiting differential methylation in either TSS, gene body, or TES regions (SData 11, SData 12).

### DEDMGs shared by all ecotypes exhibited similar regulation

DEDMGs shared in all ecotypes were identified (Figure 4E, Fig. S16A, SData 13). Thirteen of the fourteen identified in crowns (Figure 4E) exhibited the same direction of expression change in all ecotypes (either both upregulated or downregulated for each gene). Most of the downregulated DEDMGs were methylated at CHH sites within gene bodies (SData 13). In crowns, examples of genes with similar regulation included one encoding phytochrome E (*FvH4_3g06380*), one for a Cysteine-rich secretory protein, Antigen 5, and Pathogenesis-related 1 protein superfamily protein (*CAP*, *FvH4_4g09940*), both downregulated and methylated in TSS regions, whereas an *FAD/NAD(P)-*binding oxidoreductase family protein-encoding gene (*FvH4_1g23740*) was upregulated and methylated in gene bodies. A *MADS*-box transcription factor family protein-encoding gene (*FvH4_5g35401*) was upregulated in crowns of ‘Alta’ and FDP817’ and hypomethylated at CHG and CHH sites in ‘Alta,’ and at CHH in ‘FDP817’ and ‘NCGR1363’ within gene bodies of crowns in all three ecotypes.

### DEDMGs shared between ecotypes exhibited similar regulation and methylation patterns across regions and contexts

Next, we analyzed DEDMGs shared by pairs of ecotypes. ‘Alta’ shared more DEDMGs with ‘FDP817’ (crowns, 113; leaves, 107) and ‘NCGR1363’ (crowns, 104; leaves, 112) than ‘FDP817’ shared with ‘NCGR1363’ (crowns, 20; leaves, 39) (Fig 4E; SData 12). Over 90 % of shared DEDMGs were differentially expressed in the same directions in ecotypes (AC∩FC, 91 %; AC∩NC, 92 %; FC∩NC, 100 %; AL∩NL, 97 %; AL∩FL, 100 %; FL∩NL, 97 %), with most of these methylated in the CHH context (SData 14).

Among shared DEDMGs, we identified those exhibiting similar expression changes between ecotypes (Figure 5A). Each gene in 5A was upregulated or downregulated in both ecotypes (Figure 5B). Within each expression group, a gene was hyper- and/or hypomethylated in both ecotypes (Figure 5C) and in one or multiple genic regions (Figure 5, panels D-F; numbers in boxes). Twenty to 37 % of genes in column C showed similar methylation in both ecotypes: each gene in this category was either hypermethylated or hypomethylated in one or multiple genic regions (Fig. 5D-F; Fig. S17, SData 14).

**Figure 5.**
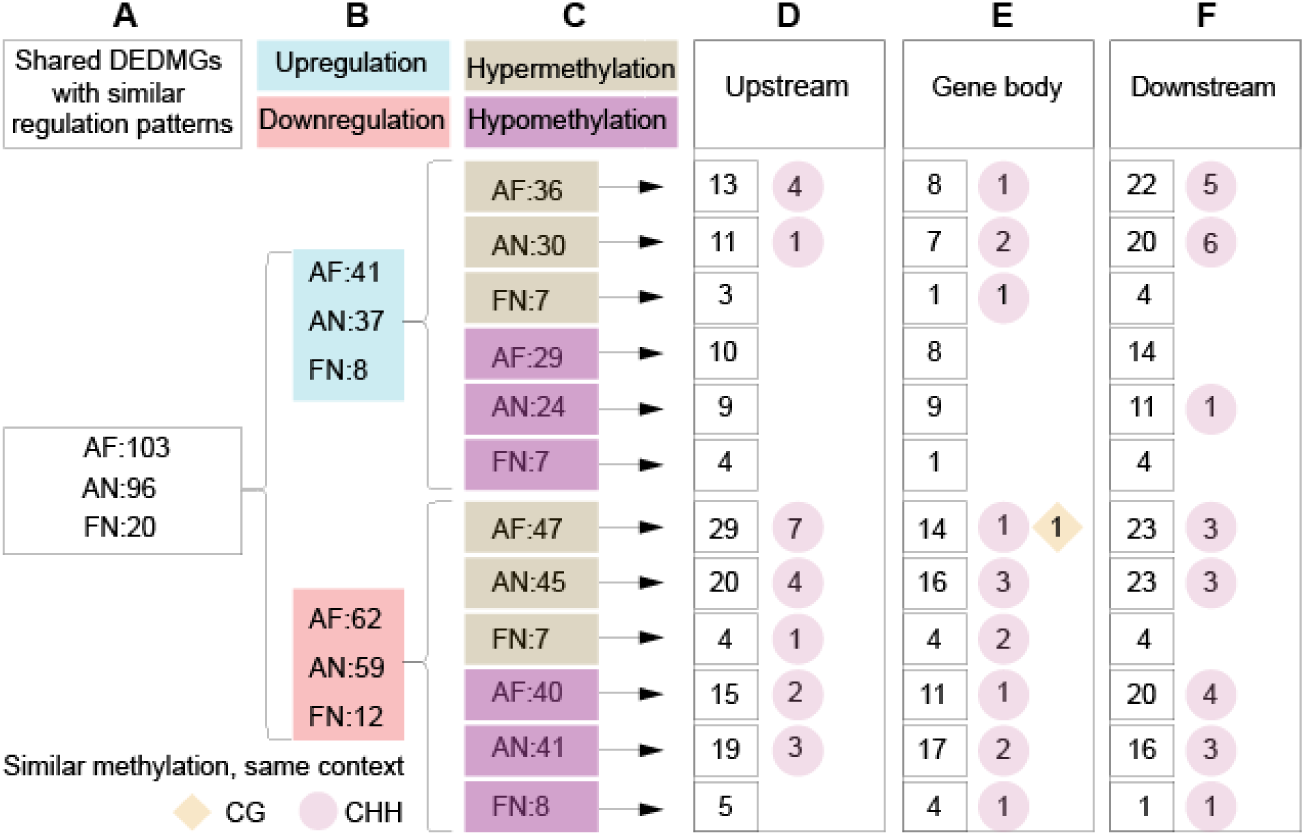
Analysis of crown DEDMGs shared between ecotypes. (A) Numbers of shared DEDMGs differentially expressed in the same directions between ecotypes. (B) Subsets of shared DEDMGs expressed in the same directions. (C) DEDMGs from column B with differential methylation in the same directions. (D-F) Numbers of shared DEDMGs from column C, showing differential methylation in the same directions in genic regions (squares) and contexts within regions (diamond and circles) of ecotype pairs. The diamond and circles represent CG and CHH methylation contexts, respectively. AF= ‘Alta’ and ‘FDP817,’ AN= ‘Alta’ and ‘NCGR1363,’ FN= ‘FDP817’ and ‘NCGR1363.’

In crowns of ‘Alta’ and ‘FDP817,’ a *HEAT SHOCK COGNATE PROTEIN 70-1*-encoding gene (*FvH4_7g22332*) was downregulated and hypermethylated in CHH sites of the TSS region. A gene encoding a leucine-rich repeat receptor-like protein kinase family protein (*FvH4_5g37590*) was also identified, with approximately 9 times higher expression in ‘Alta’ than in ‘FDP817’ and CHH hypermethylation in TES regions. Crowns of ‘Alta’ and ‘NCGR1363’ shared *CYP71B34* (*FvH4_4g25390*) and *GLUTATHIONE PEROXIDASE 6* (*FvH4_5g02130*), both hypermethylated at CHH sites in their TES regions and upregulated in ‘Alta.’ Both genes showed approximately 43-fold and three-fold higher expression, respectively, in cold-treated ‘Alta’ compared to the corresponding response in ‘NCGR1363.’ In crowns of ‘FDP817’ and ‘NCGR1363,’ an F-box family protein encoding gene (*FvH4_3g27720*) was upregulated and hypermethylated in gene bodies, while *PLAT/LH2 DOMAIN-CONTAINING LIPOXYGENASE FAMILY PROTEIN* (*FvH4_4g15250*) and *AGAMOUS-LIKE 16* (*FvH4_5g34080*) were both downregulated and hypermethylated at CHH sites (SData 14). In leaves, ‘Alta’ and ‘FDP817’ shared the DEDMG *HISTONE H3 K4-SPECIFIC METHYLTRANSFERASE SET7/9 FAMILY PROTEIN* (*FvH4_5g20710*), an upregulated gene with a 10-fold higher expression in ‘FDP817’ than in ‘Alta,’ hypermethylated at CG sites in gene bodies. Examples of genes with similar patterns shared by leaves of ‘FDP817’ and ‘NCGR1363’ include ethylene-responsive transcription factor-encoding gene *RAP2-4* (*FvH4_1g21210*), downregulated and hypomethylated at CHH sites in either TSS or TES regions, and *MINICHROMOSOME MAINTENANCE 9* (*MCM9*; *FvH4_2g10130*), upregulated and hypomethylated at CHH sites in the upstream region (SData 14).

We also identified shared DEDMGs with the same differential methylation directions that exhibited incongruent differential expression directions, and those with the same differential expression directions but contrasting differential methylation directions (SData 15 and 16). To further elucidate potentially universal *Fragaria* responses to low-temperature stress given our experimental design, these incongruent categories of shared DEDMGs are reported in the supplementary materials.

### Crowns and leaves exhibited varied DNA methylation and transcript abundance responses to cold acclimation

Expression and methylation profiles of DEDMGs in crowns and leaves within each ecotype were examined to assess molecular dynamics that may contribute to differences in tissue susceptibility to cold stress. Crowns and leaves of ‘Alta,’ ‘FDP817’ and ‘NCGR1363’ shared 235 (19 %), 49 (8 %) and 39 (8 %) DEDMGs, respectively. In all three ecotypes, more than 80 % of shared DEDMGs were either up- or downregulated in both tissues, 50 % in ‘Alta’ were either hypermethylated and/or hypomethylated in both tissues in the same genic regions and contexts, with similar patterns observed for 32 % and 24 % of shared DEDMGs in ‘FDP817’ and ‘NCGR1363,’ respectively (Fig. S18, SData 14). Among genes in this category, we identified genes with known and lesser-known roles in plant responses to low-temperature stress. In ‘Alta,’ *DEHYDRIN2-like* (*FvH4_1g12582*) was hypomethylated at CHH sites in the gene body and upregulated with fold change values of 64 and 134 in crowns and leaves, respectively. Other upregulated genes included *COLD-REGULATED 413-PLASMA MEMBRANE 2* (*COR413-PM2; FvH4_2g07440*), *GIBBERELLIN-REGULATED PROTEIN 1* (*FvH4_2g38480*), and *CYP71A25* (*FvH4_7g13060*). In ‘FDP817,’ genes encoding a late embryogenesis abundant (LEA) hydroxyproline-rich glycoprotein family (*FvH4_6g20140*) and an alpha/beta-hydrolases superfamily protein (*FvH4_2g33160*) were identified as upregulated, while in ‘NCGR1363,’ an upregulated gene encoding a photosystem II 11 kDa protein-related protein (*FvH4_4g15520*) was identified in crowns and leaves of ‘NCGR1363.’ Additional genes with similar expression and methylation patterns included *SERINE CARBOXYPEPTIDASE-LIKE 7* (*FvH4_2g03522*) and *GLYCOSYL TRANSFERASE FAMILY 2* (*FvH4_6g17730*) in ‘Alta,’ *LACCASE 14 (FvH4_1g20010)* and *UDP-GLUCOSYL TRANSFERASE 85A3* (*UGT85A3*; *FvH4_3g17550*) in ‘FDP817,’ and *SEC7 DOMAIN-CONTAINING PROTEIN* (*FvH4_7g07780*) in ‘NCGR1363,’ all downregulated in both tissues of their respective ecotypes. Twenty-nine (35 %) of the shared DEDMGs with similar regulation exhibited opposite methylation patterns between crowns and leaves in ‘Alta,’ 13 (57 %) in ‘FDP817’, and eight (47 %) in ‘NCGR1363’ (SData 14); among these were known examples of cold-responsive genes.

## Discussion

This study elucidates the influence of cold stress on DNA methylation and gene expression in two main tissues of three *F. vesca* ecotypes exhibiting contrasting cold tolerances. Collectively, our analyses reveal cold temperature-induced epigenetic dynamics and associated shifts in steady-state transcript abundance that differentiate woodland strawberry ecotypes of contrasting cold hardiness.

### The most cold-adapted ecotype exhibited enhanced epigenetic plasticity

Methylation levels varied among the three ecotypes in both control and cold-treated samples, suggesting that population history influences methylation dynamics. In addition, ‘Alta’ exhibited higher methylation levels both at the basal level in control plants and after cold treatment compared with ‘FDP817’ and ‘NCGR1363.’ The ability of *Fragaria vesca* to adapt to its local environment, including responses to temperature changes, is likely influenced by DNA methylation (Sammarco et al. 2022). Therefore, the higher epigenetic plasticity exhibited by ‘Alta’ compared with ‘FDP817’ and ‘NCGR1363’ in response to low-temperature stress may contribute to its enhanced cold-hardy phenotype.

CG methylation (mCG) was prevalent in gene bodies in our studies, consistent with genome-wide patterns reported for woodland strawberry under temperature stress (López et al. 2022) and winter rapeseed under freezing stress (Zheng et al. 2022). CG methylation was higher in all three genic regions, with a sharp decline at the transition regions between them. Low prevalence of mCG in and around TSS and TES regions is suggested to be crucial for gene expression due to possible inhibition of RNA polymerase II (Pol II) initiation by mCG (Zilberman et al. 2007; Zemach et al. 2010). DNA methylation changes were more common in CHG and CHH contexts after cold treatment. Although the CHH context showed the lowest methylation levels, consistent with previous reports in maize (Li et al. 2015), cold-induced changes in CHG and CHH methylation indicated sensitivity of non-CG methylation to environmental stress (Sammarco et al. 2024). Studies have suggested that *cis*-regulatory elements found in the promoter and enhancer regions of a gene are frequently derived from TEs (Lisch 2013). Loss of DNA methylation in non-CG contexts has been associated with loss of TE inhibition, leading to defects in plant growth and development (Cheng et al. 2015; Hu et al. 2021). In the current study, the average number of DMRs was approximately nine times higher in TEs than in genic regions, consistent with the view that most methylation responses under stress occur in transposons (Peña-Ponton et al. 2024). The abundance of TE methylation observed before and after cold acclimation further underscores the relevance of TE silencing in maintaining genome stability and suppressing cryptic TSS, whose activation changes the regular transcription activity of proximal *bona fide* TSS (Talarico et al. 2024; Le et al. 2020). Our data also showed that differential methylation was more frequent at CHH sites than in other contexts, suggesting that DNA methylation in these ecotypes was predominantly mediated by genes encoding *DRM1/DRM2* and *REPRESSOR OF SILENCING 1 (ROS1) DNA* glycosylases (Erdmann and Picard 2020; Liu et al. 2023).

### Ecotypes mobilize different cold-responsive pathways during cold acclimation

Our RNA-seq studies revealed more extensive cold-induced gene regulation in ‘Alta,’ with stronger responses in crowns than in leaves, and with more than 50 % of DEGs downregulated. The enhanced response to cold in crowns may reflect the high susceptibility of their large pith cells to cold-induced injury (Zareei et al. 2021). Plants respond to cold stress by regulating pathways that maintain metabolic balance to preserve cellular homeostasis, sustain energy production, and enhance survival under low temperature conditions. GO enrichment revealed DEDMGs in pathways that were ecotype-specific as well as shared with CHH-context–specific responses, including processes essential for cold acclimation.

Among the GO terms uniquely enriched in each ecotype were several associated with established plant stress-acclimation strategies. In ‘Alta’, crowns were enriched for ‘cellular carbohydrate metabolic process’ and ‘ATP hydrolysis activity’, consistent with maintenance of energy supply (Yu et al. 2020), whereas leaves were enriched for ‘cell redox homeostasis’ and ‘photosystem,’ consistent with ROS detoxification and protection of photosynthetic function (Hajihashemi et al. 2018; Herath 2018). In ‘FDP817,’ crowns were enriched for ‘microtubule binding,’ and leaves for ‘glycerol ether metabolic process’ and ‘alpha-amino acid metabolic process,’ suggesting cytoskeletal and metabolic adjustments during cold acclimation (Jiang et al. 2022; L. Wang et al. 2019; Moellering et al. 2010). These processes have been associated with membrane stability and maintenance of cellular homeostasis during cold stress. In ‘NCGR1363,’ crowns were enriched for ‘systemic acquired resistance,’ a characteristic of distal defense signaling (Miller et al. 2009), whereas leaves were enriched for ‘1,4-alpha-glucan branching enzyme activity,’ which is associated with branch formation in amylopectin during starch metabolism (Azeem et al. 2016).

Some enriched GO terms were identified in more than one ecotype/tissue; ‘Heme binding’ and ‘glycosyltransferase activity’ were shared by ‘Alta’ crowns and leaves. DEDMGs identified in the ‘heme binding’ pathway are involved in oxidoreduction processes essential for eliminating cold-induced ROS in plants (Balestrasse et al. 2010; Sehrawat et al. 2013). Glycosyltransferases catalyze glycosylation, leading to the accumulation of flavonoids, secondary metabolites whose cold-induced increase is associated with low temperature tolerance (Hannah et al. 2006). Leaves of ‘Alta’ and ‘FDP817’ shared ‘protein-disulfide reductase.’ (Park et al. 2021) reported that the disulfide reductase activity of the antioxidant THEOREDOXIN H2, also identified in this study, contributed to cold stress tolerance in *A. thaliana*.

Half of the pathways showing enrichment in DEDMGs overlapped with GO terms enriched in DEGs. Among the shared pathways with important roles in cold stress tolerance were carbohydrate metabolic processes, ATP metabolic processes, cellular homeostasis, and detoxification. Among GO terms enriched only in DEGs, we identified ‘catechol oxidase activity.’ *CATECHOL OXIDASE* is a polyphenol oxidase whose increased accumulation catalyzes oxidation and polymerization of phenolic compounds, secondary metabolites suggested to contribute to cold stress tolerance in grapevine (Król et al. 2015). This finding highlights the critical role of genetic and epigenetic regulation in plant cold response and tolerance. Shared pathways identified in the enrichment analysis may indicate common regulatory mechanisms among ecotypes, whereas unique pathways indicate diversity in cold-response processes that may underlie ecotype differences in cold tolerance.

### Cold-induced hyper- and hypomethylation showed no clear association with differential gene expression

Analysis of the relationship between methylation patterns and transcript accumulation levels showed that 8-17 % of DMGs also exhibited differential expression (Figure 4A, Fig S14A), indicating that most DMRs overlapping genic regions were associated with genes whose expression was not regulated by cold stress. Similar patterns have been reported in drought-and heat-stressed strawberry (Cao et al. 2022; Zhang et al. 2023), cold-stressed rapeseed (Zheng et al. 2022), and grafted rubber tree (Li et al. 2024). In contrast, a higher proportion of DEDMGs was reported in rice under cadmium stress. (Feng et al. 2016).

Analysis of the relationship between methylation patterns and transcript accumulation levels also showed a similar distribution of hyper- and hypomethylated regions between up- and downregulated genes (López et al. 2022), although a negative correlation between methylation and expression has been reported in drought-stressed rice (Wang et al. 2016). High pigmentation was observed in sweet orange fruit exhibiting increased expression of anthocyanin biosynthesis-related genes, whose promoters were hypomethylated during cold stress (Sicilia et al. 2020).

Findings from this study confirm previous reports that gene expression is not affected solely by DNA methylation (Lucero et al. 2020; Gusain et al. 2023). The relationship between methylation direction in CG sites upstream of genic regions and gene regulation revealed that at least 50 % of hypermethylated genes were downregulated in crowns of ‘Alta’ and leaves of ‘Alta,’ ‘FDP817’, and ‘NCGR1363.’ Our data are in agreement with reports that promoter methylation is most likely associated with gene repression under abiotic stress in plants (Al-Harrasi et al. 2018; Bilichak et al. 2012) and that genes methylated at promoters tend to be expressed in a tissue-specific manner (Zhang et al. 2006). While some authors suggest that gene body methylation enhances transcription (Zilberman et al. 2007), others report that the effect of gene body methylation on gene expression is specific to the tissue, ecotype, or context (Al-Harrasi et al. 2018; Wang et al. 2016) in stressed plants, which is in line with our findings.

### Cold acclimation induced different responses in ‘Alta,’ ‘FDP817’ and ‘NCGR1363’

This study demonstrated that ‘Alta’, which is adapted to the lowest median winter temperature of the three ecotypes analyzed, exhibits the greatest genetic and epigenetic plasticity in response to cold acclimation. ‘Alta’ originates from high latitude (69.9° N) in Northern Norway, which is geographically distant and climatically different from the warm California climate where ‘FDP817’ was sampled (34.4° N). The ‘NCGR1363’ ecotype originates from Bolivia, located near 20° S latitude. The different climatic conditions associated with these sites of origin likely contribute to variability in adaptation thresholds associated with cold tolerance in these ecotypes. The differing cold tolerance phenotype exhibited is suggestive of different pathways being employed during cold acclimation, as also exemplified in metabolomic studies in *Fragaria* (Davik et al. 2013; Rohloff et al. 2012).

Each ecotype responded to cold through the mobilization of its own set of DEGs, DMGs, and DEDMGs. This study reveals that the universal cold response in *Fragaria vesca*, in terms of differential gene expression, cannot be mechanistically attributed to differential DNA methylation. Only a few DEDMGs were shared by all three ecotypes, indicating a common, universal *Fragaria* response to cold stress. Among these DEDMGs were genes encoding cold stress-related transcription factor family proteins, and proteins involved in pathogenesis resistance, cellular detoxification, and signaling. For example, cold induction of *MADS*-box transcription factor family genes in pepper increased plant survival and reduced cold-induced ROS accumulation during acclimation (R. Chen et al. 2019). Similarly, in apple, expression of *MADS*-box genes during winter promotes growth cessation and bud dormancy, whereas their silencing in spring is required for dormancy release and subsequent growth resumption. (Moser et al. 2020). We identified *PHYTOCHROME E*, *FAD/NAD(P)-*binding oxidoreductase family protein and *CYTOCHROME P450* among genes associated with cold stress response. These genes have been reported to function as ROS scavengers, and their activities are associated with enhanced cold stress adaptation in plants (Qiu et al. 2023; Chakraborty et al. 2023; Tang et al. 2021). Zinc finger motif-containing proteins have been associated with increased cold tolerance (Han et al. 2021); a transgenic tobacco overexpressing a cotton *GhZFP1* gene (*ZINC FINGER C-X8-C-X5-C-X3-H TYPE*) showed increased resistance to high salt and fungal disease, and *GhZFP1* gene is drought inducible in cotton (Guo et al. 2009). In finger lime, a *CAP* gene (encoding a Cysteine-rich secretory protein, antigen 5, and pathogenesis-related 1 protein) showed increased transcript levels in response to cold stress in leaves, and high *CAP2* expression was associated with pathogen resistance and with mitigating oxidative stress responses (Mahmoud et al. 2024). PHYTOCHROME E is among the photoreceptors involved in light signalling for the activation of cold-induced genes such as CBFs through C/DRE (C-repeat/dehydration-responsive element) in cold-stressed plants (Kim et al. 2002). Cold-induced DNA hypomethylation in promoters of an *OST1* homolog in rice, as well as *ICE1* and *CBF2* in rubber plants, has been associated with upregulation of these genes and increases in cold tolerance (Guo et al. 2019; Tang et al. 2018).

Although ‘Alta,’ ‘FDP817,’ and ‘NCGR1363’ have diverse origins and cold tolerance levels, pairwise comparisons revealed shared DEDMGs among ecotypes, indicating common regulatory mechanisms. Analysis of each ecotype pair provided potentially relevant explanations for shared features. In crowns, ‘Alta’ shared more DEDMGs with ‘FDP817’ than with ‘NCGR1363.’ Given the extreme differences in cold tolerance between ‘Alta’ and ‘FDP817,’ it is notable that their DEDMG profiles were more similar than those of ‘Alta’ and ‘NCGR1363.’ Shared DEDMGs between ‘Alta’ and ‘FDP817’ could potentially reflect common characteristics of northern hemisphere ecotypes, whereas ‘NCGR1363’ originates from the southern hemisphere. Among DEDMGs shared by ‘Alta’ and ‘FDP817’ was a gene encoding a HISTONE H3 K4-SPECIFIC METHYLTRANSFERASE SET7/9 FAMILY PROTEIN, whose involvement in mitigating ROS induced by oxidative stress has been reported in animals (Daks et al. 2023). In *A. thaliana*, activation of histone lysine (K) methyltransferase (HKMTases) with SET domains in response to cold triggers the timely induction of *VIN3* (*VERNALIZATION INSENSITIVE 3*), whose expression is critical for *FLC* (*FLORAL LOCUS C*) repression during vernalization (Lee et al. 2015; Sung and Amasino 2004). *VIN3* encodes a cold-specific component that associates with PRC2 (Polycomb Repressive Complex 2) to mediate silencing of *FLC* during vernalization (Wood et al. 2006). Dehydrins are LEA (late embryogenesis abundant) proteins of the group 2 family, whose induction under cold and dehydration stress is associated with protection from cold-induced structural damage (chaperones), cryoprotection, and ROS scavenging has been reported (Szlachtowska and Rurek 2023; Close 1997). A positive correlation between H3K4me3 histone modification and increases in dehydrin expression during drought stress responses has been reported in rice (Zong et al. 2020).

‘Alta’ and ‘NCGR1363’ shared DEDMGs despite their origin from opposite sides of the equator. Based on LT_50_ values, and compared with ‘FDP817’ (LT_50_ −7,7), ‘NCGR1363’ (LT_50_-8.2) displays a hardiness level closer to ‘Alta’ (LT_50_ −11,6), possibly due to their higher altitude locations compared with ‘FDP817.’ Among shared genes were those encoding GLUTATHIONE PEROXIDASE, CYTOCHROME P450, and FAD/NAD(P)-BINDING OXIDOREDUCTASE FAMILY PROTEIN, all involved in detoxification during cold stress. Genes encoding coldLinduced transcription regulators (such as *bHLH93, MYB46, and AP2/B3*), signal transducers, and stressLresponsive genes (such as *COR413PM2* [Cold-regulated 413-plasma membrane], *LEA,* and *PHOSPHOLIPASE A 2A*) were also identified. These genes have been reported to contribute to membrane stability (Hincha and Thalhammer 2012; Zhang et al. 2021; Zhou et al. 2018), stress signaling (Ali et al. 2022; Narusaka et al. 2003), accumulation of antioxidant enzymes and CBFs (Hannah et al. 2006; Zhang et al. 2024), regulation of antioxidant activity (Pei et al. 2024; Yang et al. 2023), activation of metabolite biosynthesis and secondary cell wall biosynthesis pathways (K. Chen et al. 2019), and promotion of vernalization-induced flowering (Sharma 2011).

‘FDP817’ and ‘NCGR1363’ have the closest and least extreme LT_50_ values and are both located near the equator, which may help explain the DEDMGs shared between these ecotypes. Among the downregulated DEDMGs was *AGAMOUS-LIKE 16* (*AGL16*), a MADS-box transcription factor that negatively regulates abiotic stress responses (Zhao et al. 2021; Lei et al. 2024) and flowering time by repressing *FLOWERING LOCUS T* (*FT*) (Hu et al. 2014). Among the upregulated DEDMGs were a *DEAD/DEAH-BOX RNA HELICASE FAMILY PROTEIN* gene and *MINICHROMOSOME MAINTENANCE 9* (*MCM9*). *MCM9* is a DNA helicase involved in maintaining DNA integrity and genome stability in animals by protecting the DNA replication fork from degradation following stress-induced stalling (Griffin et al. 2022). Upregulation of *MCM9* in ‘Alta’ and ‘FDP817’ suggests its crucial role in preventing DNA damage in plants under cold stress. A similar pattern was exhibited by an *MCM9* homolog in cold-tolerant peach varieties (Nilo-Poyanco et al. 2019). Cold induction of the *DEAD-BOX RNA HELICASE*-encoding gene was associated with cold tolerance through enhancement of CBF gene expression in *A. thaliana* (Gong et al. 2002). The repressive role of DEAD-box RNA helicases on stress-responsive transcription factors via the *RdDM* pathway has been reported (Barak et al. 2014).

Analysis of transcript and methylation levels of shared DEDMGs reveals that each ecotype exhibits unique stress response regulation patterns, likely related to their chilling requirements. These differences may result from accumulated mutations during evolution and adaptation. For example, in rice ecotypes, distinct methylation patterns in DEDMGs with similar expression profiles were associated with differentially methylated cytosine SNPs (single nucleotide polymorphisms; (Rajkumar et al. 2020).

Differences in cold tolerances among ‘Alta’, ‘FDP817’, and ‘NCGR1363’ may reflect accumulated genetic and epigenetic variation, arising either randomly or in response to environmental cues. In this study, more than 65 % of identified DEDMGs were ecotype-specific. These unique DEDMGs included stress-induced genes involved in stress tolerance, transcriptional regulation, carbohydrate metabolism, cell-membrane stability, and oxidative stress reduction. In ‘Alta,’ genes encoding *DEHYDRATION-RESPONSIVE ELEMENT-BINDING PROTEIN 1E* (*DREB1E*), *WRKY*, *MADS-BOX TRANSCRIPTION FACTOR FAMILY PROTEIN,* and *HEAT SHOCK TRANSCRIPTION FACTOR C1* were identified in crowns, while a *XYLOGLUCAN ENDOTRANSGLYCOSYLASE, CELLULOSE SYNTHASE* involved in carbohydrate metabolic processes, was regulated in leaves. Roles of transcription factors and their diverse expression patterns in relation to cold tolerance have been reported in plants (Zhang et al. 2019). Among DEDMGs unique to ‘FDP817’ were genes encoding RING-H2 finger proteins such as ATL20 and ATL7, whose role in enhancing stress tolerance has been reported in apricot kernel and grapevine (Liu et al. 2022; L. Wang et al. 2017). ‘FDP817’-specific DEDMGs also included those encoding endochitinases, which function as antifreeze proteins and contribute to pathogen resistance (Griffith and Yaish 2004). For ‘NCGR1363,’ unique DEDMGs included *SUMO-CONJUGATING ENZYME 1* (*SCE1*), whose overexpression enhances drought and salt tolerance in transgenic tobacco and *Arabidopsis thaliana* (H. Wang et al. 2019b, 2019a) but reduces drought tolerance in transgenic rice (Joo et al., 2019). Additionally, *EARLY RESPONSIVE TO DEHYDRATION 15*, encoding transcription factor *ERD15*, was identified; its cold-induced expression enhances cold tolerance in grapevine (Yu et al. 2017).

### *F. vesca* crowns and leaves have distinct cold response mechanisms

Results presented in this study show that eight to 19 % of DEDMGs identified in ecotypes were tissue-specific. More than 80 % of these were regulated in the same direction in all respective ecotypes, and less than half of them exhibited similar methylation patterns. This finding suggests that tissues within ecotypes can employ distinct methylation and expression mechanisms in response to stress. Similarly, tissue-specific methylation and expression have been reported in wheat and rice ecotypes exposed to salt stress (Kumar et al. 2017; Ratna et al. 2012). Among DEDMGs shared by crowns and leaves of each ecotype, we identified *MYO-INOSITOL OXYGENASE 1* and *FLAVONOL SYNTHASE 1* in ‘Alta,’ *LACCASE 14* and *EXPANSIN A4-LIKE* (*EXLA4*) in ‘FDP817’, and *SEC7 DOMAIN-CONTAINING PROTEIN* in ‘NCGR1363.’ Myo-inositol is a sugar alcohol whose cold-induced accumulation has been associated with increased resistance to cold stress in transgenic tobacco, tomato, and rapeseed (Sambe et al. 2015; Yan et al. 2022; Wang et al. 2022; Liu et al. 2022). Laccases are critical for the polymerization of lignin, a complex polymer critical for increasing plant vigor and enhancing biotic and abiotic stress resistance (Hashemipetroudi et al. 2023). Although cold induction of laccases has been reported in plants (Zhu et al. 2023; Xu et al. 2019), their association with cold stress tolerance has not been established. Expansins are proteins involved in loosening and extension of the cell wall (Cosgrove 2000), with *expansin A4* mutant lines reported to exhibit hypersensitivity to salt and drought stress in plants (Chen et al. 2018). ‘NCGR1363’ crowns and leaves shared a *SEC7 DOMAIN-CONTAINING PROTEIN*, previously shown to be downregulated during cold and drought stress, with overexpressing lines exhibiting cold tolerance (López-Cristoffanini et al. 2015; Ashraf and Rahman 2019).

## Conclusion

This study integrates investigation of DNA methylation and gene expression to identify candidate genes and pathways underlying differences in cold tolerance in ‘Alta,’ ‘NCGR1363’ and ‘FDP817.’ Crowns and leaves of these three ecotypes of *F. vesca* exhibited distinct cold response mechanisms. A majority (>65 %) of DEDMGs were ecotype-specific. Despite the diverse origins and cold tolerances of the ecotypes studied, pairwise comparisons revealed shared DEDMGs and cold response pathways between them, indicating common regulatory mechanisms. Correlation between cold-induced methylation and transcript accumulation revealed that cold responses in *Fragaria vesca*, as reflected in gene expression, cannot be mechanistically attributed to DNA methylation. ‘Alta,’ the most cold-tolerant of the three analyzed ecotypes, exhibited the greatest genetic and epigenetic plasticity in response to low-temperature acclimation and also showed higher basal methylation levels in tissues and contexts, compared to ‘NCGR1363’ and ‘FDP817.’ Global methylation analysis showed changes in DNA methylation, especially in CHG and CHH contexts, following cold treatment. CG methylation was higher in gene bodies, while non-CG methylation was enriched in regions upstream of TSS and downstream of TES regions. A majority of cold-induced DMRs were hypomethylated in crowns of ‘Alta,’ whereas most DMRs in ‘NCGR1363’ and ‘FDP817’were hypermethylated.

Ecotype-specific expression and methylation patterns in ‘Alta,’ ‘FDP817’ and ‘NCGR1363’ may be crucial for local adaptation to low temperatures, a quantitative trait influenced by multiple genes. Variation in methylation and expression patterns identified in this study indicates that cold signaling and survival depend on tissue type and the geographic origin of the plants exposed to cold stress. Results from this study support a potential role for epialleles, in addition to genetic markers, for use as molecular markers for crop improvement. Targeted functional analysis of the candidate markers identified in this study represents a clear future direction. Precise genome and epigenome editing approaches in diploid *Fragaria* will be particularly valuable for elucidating the complex pathways and networks underlying low temperature stress responses across the Rosaceae.

## Acknowledgements

We thank Gage Koehler, University of Indiana, for help with DNA isolation. We also thank Jose Gutierrez-Marcos and Ryan Merritt, University of Warwick, for valuable input on DMR calling parameters and data analysis. Sequencing service was provided by the Norwegian Sequencing Centre (www.sequencing.uio.no), a national technology platform hosted by the University of Oslo and Oslo University Hospital. The computational analysis was partly performed on resources provided by Sigma2 - the National Infrastructure for High-Performance Computing and Data Storage in Norway (www.sigma2.no).

## Author contribution

RCW, PEG, and SKR designed the research plan. JD generated the plant material. RGN performed the experiments. RCW, PEG, and RGN analyzed data and wrote the original draft. RCW, PEG, RGN, SKR, MA, and WJ edited the manuscript. All authors revised and approved the manuscript.

## Supplemental figure legends

**Figure S1. Global DNA methylation landscape in leaves and crowns of ‘Alta,’ ‘FDP817’, and ‘NCGR1363’ ecotypes.** (A) DNA methylation levels at CG, CHG, and CHH contexts in chromosomes (Fvb1-7) of ‘Alta,’ ‘FDP817’, and ‘NCGR1363’ ecotypes cold acclimated at 2 °C (CT) for 42 days. (B) DNA methylation levels (methylated reads/total reads) at CG, CHG, and CHH contexts in chromosomes (Fvb1-7) of ‘FDP817’ and ‘NCGR1363’ propagated at 20 ± 2 °C (UT) for 42 days. Lanes 1-6 show methylation levels for each tissue and context. Methylation levels are shown for all methylation contexts along all seven *F. vesca* chromosomes, using a 50 kb window. CT= Cold-treated, UT= Untreated.

**Figure S2. Global DNA methylation patterns in untreated leaves and crowns of ‘Alta,’ ‘FDP817’, and ‘NCGR1363’ ecotypes.** Linear plots show DNA methylation levels (methylated reads/total reads) for crowns and leaves at CG, CHG, and CHH contexts in chromosomes (Fvb1-7) of ‘Alta,’ ‘FDP817’, and ‘NCGR1363’ propagated at 20 ± 2 °C (UT) for 42 days. Methylation levels are shown for all methylation contexts along all seven *F. vesca* chromosomes, using a 50 kb window. ACUT= ‘Alta’ crown untreated; ALUT= ‘Alta’ leaf untreated; FCUT= ‘FDP817’ crown untreated; FLUT= ‘FDP817’ leaf untreated; NCUT= ‘NCGR1363’ crown untreated; NLUT= ‘NCGR1363’ leaf untreated.

**Figure S3. Global DNA methylation patterns in cold-treated leaves and crowns of Alta, FDP817, and NCGR1363 ecotypes.** Linear plots show DNA methylation levels (methylated reads/total reads) for crowns and leaves at CG, CHG, and CHH contexts in chromosomes (Fvb1-7) of ‘Alta’ ‘FDP817’ and ‘NCGR1363’ propagated under cold acclimation (2 °C) temperature conditions for 42 days. Methylation levels are shown for all methylation contexts along all seven *F. vesca* chromosomes, using a 50 kb window. ACCT= ‘Alta’ crown cold-treated; ALCT= ‘Alta’ leaf cold-treated; FCCT= ‘FDP817’ crown cold-treated; FLCT= ‘FDP817’ leaf cold-treated; NCCT= ‘NCGR1363’ crown cold-treated; NLCT= ‘NCGR1363’ leaf cold-treated.

**Figure S4. DNA methylation differences in untreated crowns of ecotypes.** ‘Alta’ has higher basal methylation levels (methylated reads/total reads) than ‘FDP817’ and ‘NCGR1363.’ Methylation levels are shown for all methylation contexts along all seven *F. vesca* chromosomes, using a 50 kb window. ACUT= ‘Alta’ crown untreated; FCUT= ‘FDP817’ crown untreated; NCUT= ‘NCGR1363’ crown untreated.

**Figure S5. DNA methylation differences in untreated leaves of ecotypes.** ‘Alta’ has higher basal methylation levels (methylated reads/total reads) than ‘FDP817’ and ‘NCGR1363.’ Methylation levels are shown for all methylation contexts along all seven *F. vesca* chromosomes, using a 50 kb window. ALUT= ‘Alta’ leaf untreated; FLUT= ‘FDP817’ leaf untreated; NLUT= ‘NCGR1363’ leaf untreated.

**Figure S6. DNA methylation differences in cold-treated crowns of ecotypes.** ‘Alta’ has higher methylation levels (methylated reads/total reads) than ‘FDP817’ and ‘NCGR1363.’ Methylation levels are shown for all methylation contexts along all seven *F. vesca* chromosomes, using a 50 kb window. ACCT= ‘Alta’ crown cold-treated; FCCT= ‘FDP817’ crown cold-treated; NCCT= ‘NCGR1363’ crown cold-treated.

**Figure S7. DNA methylation differences in cold-treated leaves of ecotypes.** ‘Alta’ has higher methylation levels (methylated reads/total reads) than ‘FDP817’ and ‘NCGR1363.’ Methylation levels are shown for all methylation contexts along all seven *F. vesca* chromosomes, using a 50 kb window. ALCT= ‘Alta’ leaf cold-treated; FLCT= ‘FDP817’ leaf cold-treated; NLCT= ‘NCGR1363’ leaf cold-treated.

**Figure S8. DNA methylation patterns in genomic regions of ‘FDP817’ and ‘NCGR1363’ propagated at normal (UT; 20 ± 2 °C) and low (CT; 2 °C) temperature conditions for 42 days.** (A) DNA methylation patterns in CG, CHG, and CHH sites of genic regions in ‘FDP817’ crowns and leaves. (B) DNA methylation patterns in CG, CHG, and CHH sites of TE regions in ‘FDP817’ crowns and leaves. (C) DNA methylation patterns in CG, CHG, and CHH sites of genic regions in ‘NCGR1363’ crowns and leaves. (D) DNA methylation patterns in CG, CHG, and CHH sites of TE regions in ‘NCGR1363’ crowns and leaves. Plots show cold-induced methylation changes in CHG and CHH contexts in both genic regions and TE regions. TE regions have higher basal methylation levels (UT) and undergo smaller methylation changes after cold acclimation compared with genic regions. Upstream and downstream regions are 1 kb in size. Each genomic feature and regions 1 kb upstream and downstream of these were divided into 20 windows, and the average methylation level per window was plotted to illustrate global DNA methylation levels. Colored lines show different combinations of tissue and temperature: FCUT= ‘FDP817’ crown Untreated, FLUT= ‘FDP817’ leaf Untreated, FCCT= ‘FDP817’ crown cold-treated, FLCT= ‘FDP817’ leaf cold-treated, NCUT= ‘NCGR1363’ crown untreated, NLUT= ‘NCGR1363’ leaf untreated, NCCT= ‘NCGR1363’ crown cold-treated, NLCT= ‘NCGR1363’ leaf cold-treated.

**Figure S9. DMR patterns in ecotype leaves propagated under normal (UT; 20 ± 2 °C) and low (CT; 2 °C) temperature conditions for 42 days.** (A) Distribution of differentially methylated regions at CG, CHG, and CHH contexts in all seven chromosomes of ecotypes. Numbers of DMRs in each tissue are indicated in parentheses. Red and blue dots denote hypermethylated and hypomethylated regions, respectively. (B) Number of DMRs overlapping with CG, CHG, and CHH sites in genic regions of ecotype leaves. (C) Number of DMRs overlapping with CG, CHG, and CHH sites in TE regions of ecotype leaves. ‘NCGR1363’ has the highest number of DMRs across genomic regions, with a majority of DMRs in the CHH context. Differential methylation in TE regions is about 10 times higher than in genic regions. Upstream and downstream regions are 1 kb in size.

**Figure S10. Overlap of differentially methylated genes (DMGs) between ‘Alta,’ ‘NCGR1363’ and ‘FDP817’ leaves.** (A) Numbers of shared and unique DMGs in ecotypes: ‘Alta’ and ‘NCGR1363’ share less than half of their DMGs; most DMGs in ‘FDP817’ are shared; ‘Alta’ and ‘NCGR1363’ share more DMGs than either shares with ‘FDP817.’ (B) Numbers of genes with DMRs in either one or multiple methylation context(s). A majority of genes are methylated exclusively in one context.

**Figure S11. Significantly enriched Gene Ontology (GO) terms among differentially methylated genes (DMGs) in different ecotypes.** Numbers of unique and shared GO terms at CG, CHG, and CHH contexts of ecotype crowns (A) and leaves (B). GO terms were mostly unique to each context and ecotype. (C) Shared and unique GO terms overlapping in gene bodies among ecotypes. Most enriched GO terms are shared by all ecotypes. AC= ‘Alta’ crown, AL= ‘Alta’ leaf, NC= ‘NCGR1363’ crown, NL= ‘NCGR1363’ leaf; FC= ‘FDP817’ crown.

**Figure S12. Homogeneity assessment and expression profile in crowns and leaves of ‘Alta’, ‘FDP817’, and ‘NCGR1363’ propagated at 20 ± 2 °C (UT) and 2 °C (CT) for 42 days.** (A) Clustering of replicates into treatment groups for crowns and leaves of ecotypes propagated at control (UT; 20 ± 2 °C) and cold acclimation (CT; 2 °C) conditions for 42 days. Principal component analysis revealed homogeneity within replicates of the same treatment group, except for ‘Alta’ crown (CT), for which one replicate was excluded from downstream analysis due to divergence from the other two replicates. (B) Volcano plots of expressed (-log_10_FDR) genes versus their degree of fold change (log_2_FC). Significantly expressed genes (p.adjust < 0.05) are marked in green, red, and grey, with red and green denoting genes with log_2_FC values greater than 1 (upregulated) and less than −1 (downregulated), respectively. Black dots indicate non-significant genes with p.adjust ≥ 0.05. Points within dashed lines define −1 ≤ log_2_FC ≤ 1. UT= Untreated, CT= Cold-treated.

**Figure S13. Correlation of differential methylation and gene expression in crowns of ecotypes.** (A) Overlap between DEGs and DMGs in ‘FDP817’ and ‘NCGR1363.’ (B) Number of DEDMGs harboring DMRs in either one or multiple genic regions. (C) Number of DEDMGs with DMRs in one or multiple methylation contexts. A majority of DEDMGs are methylated exclusively in one context. (D, E) Differentially expressed genes associated with DMRs (DEDMRs), showing a relationship between transcript accumulation (log_2_fold change) and differential methylation levels (proportion difference between cold-treated and untreated samples) in CG, CHG, and CHH contexts of gene bodies and regions 1kb upstream of transcription start sites and downstream of transcription end sites in ‘FDP817’ and ‘NCGR1363.’

**Figure S14. Correlation of differential methylation and gene expression in leaves of ecotypes.** (A) Overlap between DEGs and DMGs in ‘Alta,’ ‘FDP817’ and ‘NCGR1363.’ (B) Number of DEDMGs harboring DMRs in either one or multiple genic regions. (C) Number of DEDMGs with DMRs in one or multiple methylation contexts. (D-F) Differentially expressed genes associated with DMRs (DEDMRs), showing relationship between transcript accumulation (log2 fold change) and DNA methylation levels (DNA methylation proportion difference between 42D cold acclimated and control samples) in CG, CHG and CHH contexts of gene bodies and regions 1kb upstream of transcription start site and downstream of transcription end site in ‘Alta,’ ‘FDP817’, and ‘NCGR1363.’

**Figure S15. Gene Ontology (GO) enrichment analysis of differentially expressed and methylated genes (DEDMGs).** GO terms enriched in DEDMGs unique to tissues among ecotypes. Most enriched GO terms are ecotype-specific and methylation context. AC= ‘Alta’ crown, NC= ‘NCGR1363’ crown, AL= ‘Alta’ leaf, NL= ‘NCGR1363’ leaf.

**Figure S16. Differentially expressed and differentially methylated genes (DEDMGs) in leaves of ‘Alta,’ ‘FDP817’ and ‘NCGR1363.’** (A) Number of shared and unique DEDMGs in ecotypes regardless of methylation context and region. (B) Number of shared and unique DEDMGs in CG, CHG, and CHH contexts among ecotypes. Most DEDMGs are ecotype specific. AL= ‘Alta’ leaf, FL= ‘FDP817’ leaf, NL= ‘NCGR1363’ leaf.

**Figure S17. Analysis of leaf DEDMGs shared between ecotypes.** (A) Numbers of shared DEDMGs differentially expressed in the same directions between ecotypes. (B) Subsets of shared DEDMGs expressed in the same directions. (C) DEDMGs from column B with differential methylation in the same directions. (D-F) Numbers of shared DEDMGs from column C, showing differential methylation in the same directions in genic regions (squares) and contexts within regions (diamond and circles) of ecotype pairs. The diamonds, triangles, and circles represent CG, CHG, and CHH methylation contexts, respectively. AF= ‘Alta’ and ‘FDP817,’ AN= ‘Alta’ and ‘NCGR1363,’ FN= ‘FDP817’ and ‘NCGR1363.’

**Figure S18. Analysis of crown and leaf DEDMGs shared within ecotypes.** (A) Numbers of shared DEDMGs differentially expressed in the same directions between ecotypes. (B) Subsets of shared DEDMGs expressed in the same directions. (C) DEDMGs from column B with differential methylation in the same directions. (D-F) Numbers of shared DEDMGs from column C, showing differential methylation in the same directions in genic regions (squares) and contexts within regions (diamond and circles) of ecotype pairs. The diamonds, triangles, and circles represent CG, CHG, and CHH methylation contexts, respectively. A= ‘Alta’, F= ‘FDP817’, N= ‘NCGR1363.’

## Supplemental table legends

**Table S1**. **Genotypes of woodland strawberry used in this study.** The genotypes are collected from different regions and show different cold tolerance abilities. Estimated LT50 values indicate freezing temperatures at which 50 percent of cold-treated plants do not survive (Davik et al., 2013). NCGR = National Clonal Germplasm Repository. CFRA 371 and CFRA1363 are the NCGR accession IDs also identified as NCGR371 and ‘NCGR1363’, respectively, by Davik et al. (2013). CFRA371 was also denoted as ‘FDP817’ by the Horticultural Research International (HRI) in the UK (Hadonou et al. 2004). USDA-ARS, United States Department of Agriculture-Agricultural Research Service. m.a.s.l = meters above sea level.

**Table S2. WGBS mapping statistics showing the efficiency level of mapping sequenced reads of individual ecotypes to *Fragaria vesca* version 4 (FvH4.0.a2) genome.** Mapping efficiency varied from between 51 % to 70 % at the context and treatment level across ‘Alta’, ‘FDP817’, and ‘NCGR1363’ ecotypes, with leaves having a higher number of mapped reads than crowns. The majority of methylated cytosines were recorded in the CG context, followed by CHG and then CHH. ACUT= ‘Alta’ crown untreated, ALUT= ‘Alta’ leaf untreated, ACCT= ‘Alta’ crown cold-treated, ALCT= ‘Alta’ leaf cold-treated, FCUT= ‘FDP817’ crown untreated, FLUT= ‘FDP817’ leaf untreated, FCCT= ‘FDP817’ crown cold-treated, FLCT= ‘FDP817’ leaf cold-treated, NCUT= ‘NCGR1363’ crown untreated, NLUT= ‘NCGR1363’ leaf untreated, NCCT= ‘NCGR1363’ crown cold-treated, NLCT= ‘NCGR1363’ leaf cold-treated.

**Table S3. Number of DMRs showing methylation direction in CG, CHG, and CHH contexts of ‘Alta’, ‘FDP817’, and ‘NCGR1363’ tissues.** DMRs in CG and CHG contexts were called with a bin size of 100 bp, and CHH with 50 bp.

**Table S4. Number of DMRs in genic regions and contexts ‘Alta’, ‘FDP817’, and ‘NCGR1363’ tissues.**

**Table S5. Number of DMRs in TE regions and contexts ‘Alta’, ‘FDP817’, and ‘NCGR1363’ tissues.**

**Table S6. RNA-seq and mapping statistics of reads generated from Illumina sequencing of leaves and crowns of untreated (UT) and cold-treated (CT) ‘Alta’, ‘FDP817’, and ‘NCGR1363.’**

**Table S7. Number of DEDMGs in regions and contexts of ecotypes and probability of overlap between DEGs and DMGs in tissues.** (A) Total number of DMGs, DEGs, and DEDMGs. (B) Differentially expressed DMGs in genic regions and contexts. (C) Probability of overlap between DEGs and DMGs in ecotypes. Representation factor is the number of overlapping genes divided by the expected number of overlapping genes drawn from two independent groups. A representation factor > 1 indicates more overlap than expected of two independent groups, a representation factor < 1 indicates less overlap than expected, and a representation factor of 1 indicates that the two groups overlap by the number of genes expected for independent groups of genes.

## Supplemental data legends

**SData 1**. **Differentially methylated regions in crowns and leaves of ecotypes.**

**SData 2. Differentially methylated regions in genic regions of ecotypes.**

**SData 3. Differentially methylated regions in genic TE regions of ecotypes.**

**SData 4. Differentially methylated genes in crowns and leaves of ecotypes.**

**SData 5. GO enrichment in differentially methylated genes.**

**SData 6. Differentially expressed genes in crowns and leaves of ecotypes.**

**SData 7. GO enrichment in differentially expressed genes.**

**SData 8. Differentially expressed differentially methylated genes in crowns and leaves of ecotypes.**

**SData 9. Differentially expressed genes harboring differentially methylated regions in gene body, 5’ and 3’ regions of ecotypes.**

**SData 10. GO enrichment in differentially expressed differentially methylated genes.**

**SData 11. Differentially expressed differentially methylated genes unique to ecotype tissues. Columns E to M show the methylation status, region, and context of each gene.**

**SData 12**. **Differentially expressed differentially methylated genes shared among ecotypes.** In all sheets, columns display differential expression fold change values for each ecotype and methylation status of each gene across the three ecotypes, indicating methylation gain, loss, or both (gain-loss). NA denotes cases in which methylation data are not available.

**SData 13. Differentially expressed differentially methylated genes grouped as shared or ecotype-specific.**

**SData 14. Shared differentially expressed differentially methylated genes with similar regulation between ecotypes, and similar methylation patterns in regions and contexts.** Ecotypes are separated by underscore (_) in column A, with the left and right positions corresponding to Ecotype 1 and Ecotype 2 in columns D and F, respectively.

**SData 15**. **Shared differentially expressed differentially methylated genes with similar regulation between ecotypes, and opposite methylation patterns in regions and contexts.** Ecotypes are separated by underscore (_) in column A, with the left and right positions corresponding to Ecotype 1 and Ecotype 2 in column F, respectively.

**SData 16**. **Shared differentially expressed differentially methylated genes exhibiting different regulation patterns between ecotypes and similar methylation patterns in the same region and context.** Ecotypes are separated by underscore (_) in column A, with the left and right positions corresponding to Ecotype 1 and Ecotype 2 in columns D and E, respectively.

