## Supplementary Figures S1-S18 for "Epigenetic plasticity is associated with enhanced tolerance to low temperature stress in woodland strawberry"

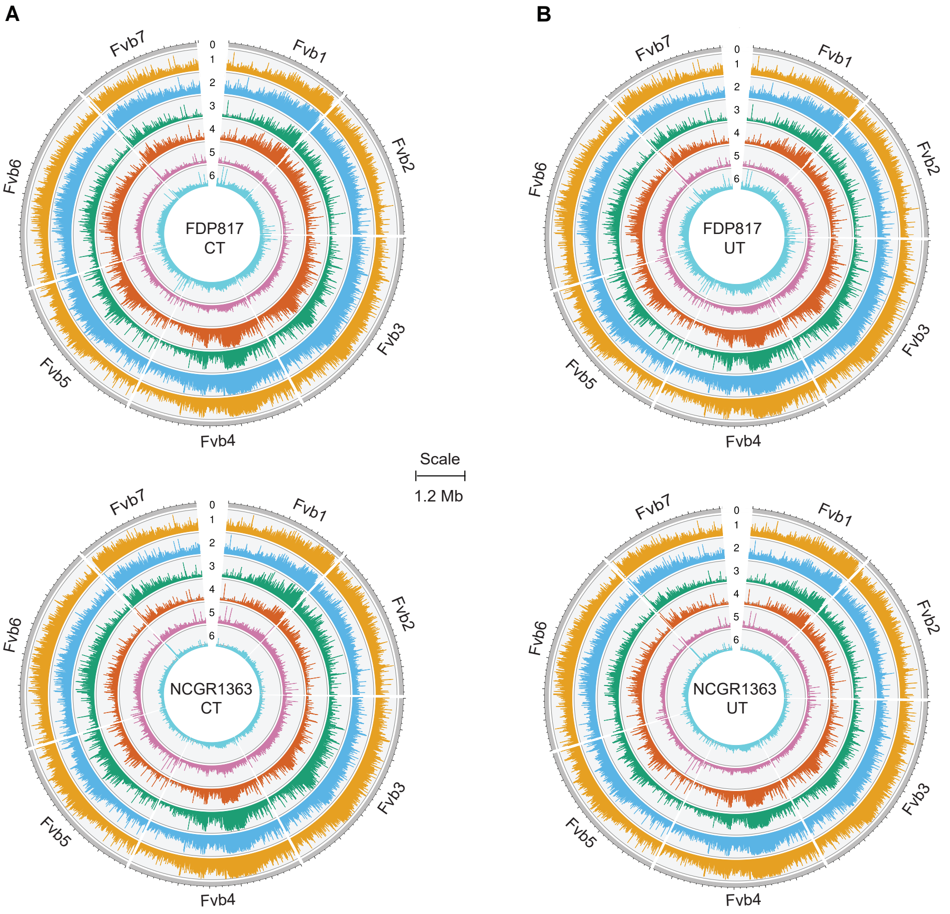


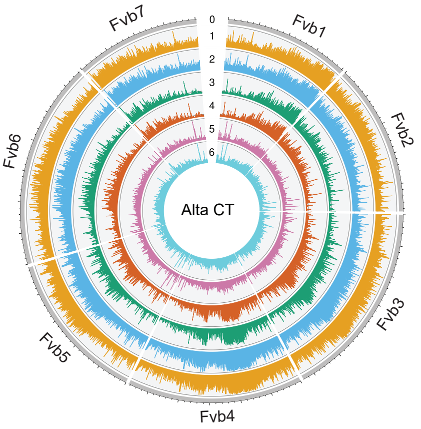


**Circos plot key**
0: Chromosome scale
1: Leaf CG context
2:Crown CG context
3: Leaf CHG context
4: Crown CHG context
5: Leaf CHH context
6: Crown CHH context

**Figure S1. Global DNA methylation landscape in leaves and crowns of 'Alta,' 'FDP817', and 'NCGR1363' ecotypes.** (A) DNA methylation levels at CG, CHG, and CHH contexts in chromosomes (Fvb1-7) of 'Alta,' 'FDP817', and 'NCGR1363' ecotypes cold acclimated at 2 °C (CT) for 42 days. (B) DNA methylation levels (methylated reads/total reads) at CG, CHG, and CHH contexts in chromosomes (Fvb1-7) of 'FDP817' and 'NCGR1363' propagated at 20 ± 2 °C (UT) for 42 days. Lanes 1-6 show methylation levels for each tissue and context. Methylation levels are shown for all methylation contexts along all seven F. vesca chromosomes, using a 50 kb window. CT= Cold-treated, UT= Untreated.

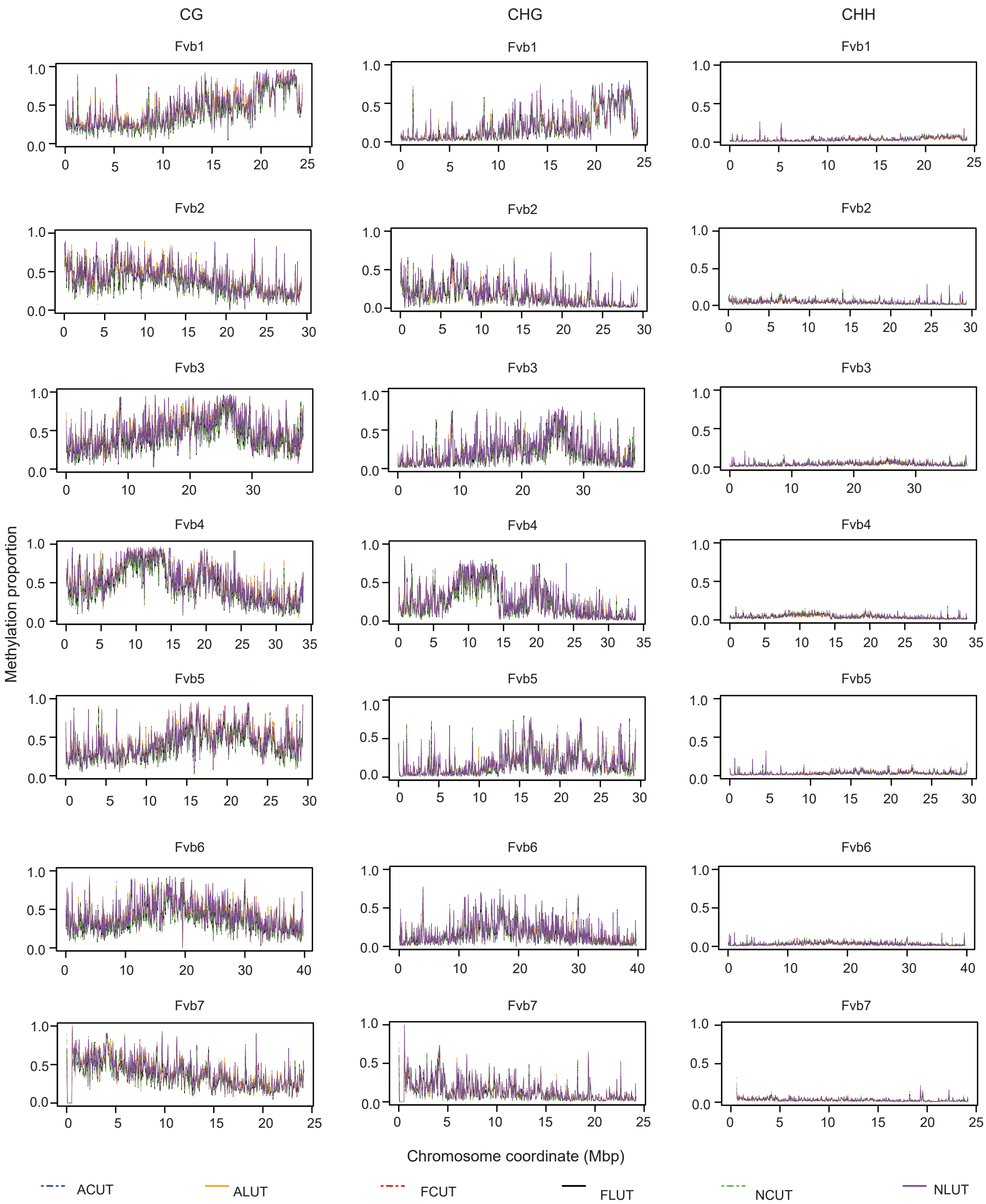


**Figure S2. Global DNA methylation patterns in untreated leaves and crowns of 'Alta,' 'FDP817', and 'NCGR1363' ecotypes.** Linear plots show DNA methylation levels (methylated reads/total reads) for crowns and leaves at CG, CHG, and CHH contexts in chromosomes (Fvb1-7) of 'Alta,' 'FDP817', and 'NCGR1363' propagated at 20 ± 2 °C (UT) for 42 days. Methylation levels are shown for all methylation contexts along all seven F. vesca chromosomes, using a 50 kb window. ACUT= 'Alta' crown untreated; ALUT= 'Alta' leaf untreated; FCUT= 'FDP817' crown untreated; FLUT= 'FDP817' leaf untreated; NCUT= 'NCGR1363' crown untreated; NLUT= 'NCGR1363' leaf untreated.

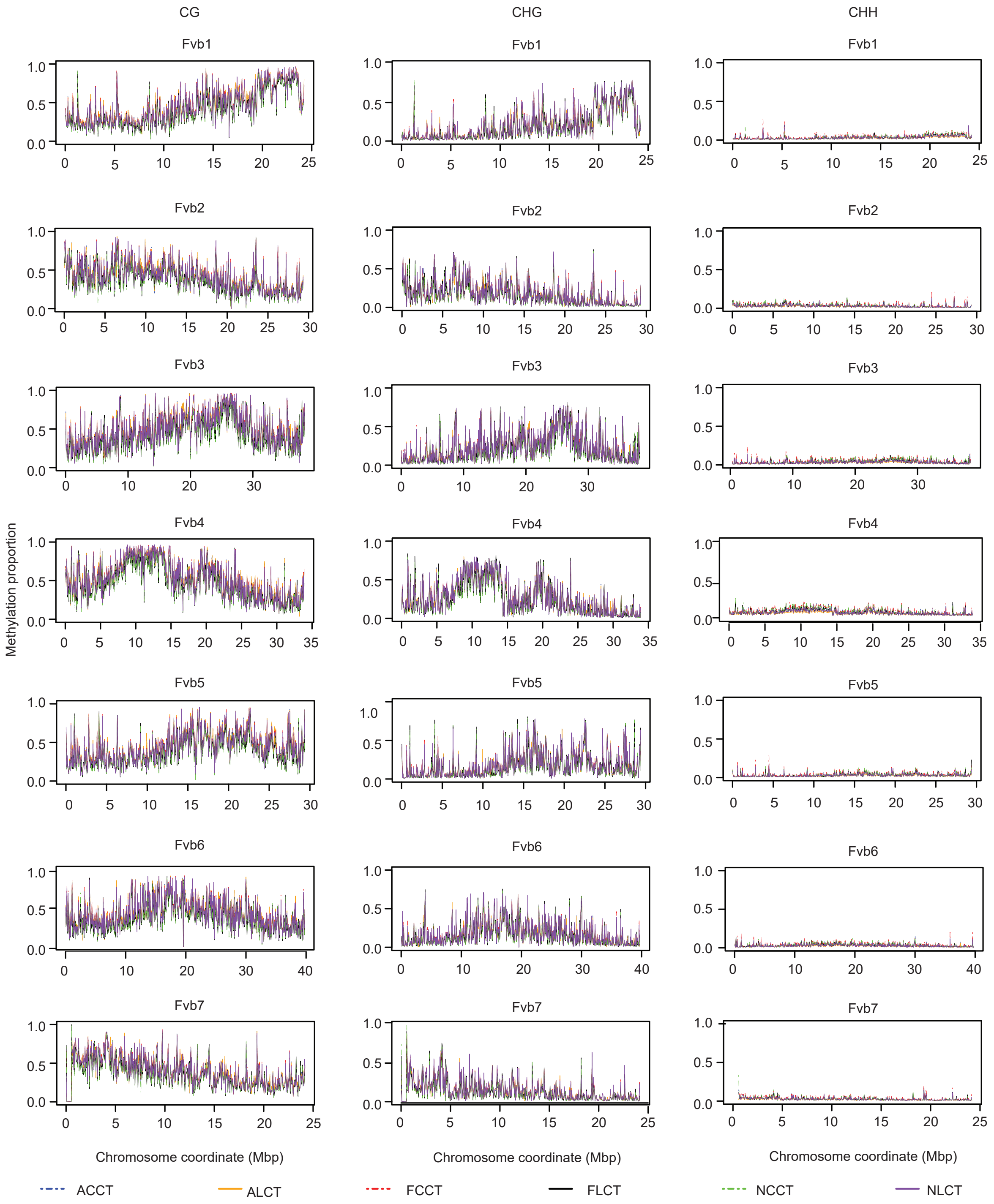


**Figure S3. Global DNA methylation patterns in cold-treated leaves and crowns of Alta, FDP817, and NCGR1363 ecotypes.** Linear plots show DNA methylation levels (methylated reads/-total reads) for crowns and leaves at CG, CHG, and CHH contexts in chromosomes (Fvb1-7) of 'Alta' 'FDP817' and 'NCGR1363' propagated under cold acclimation (2 °C) temperature conditions for 42 days. Methylation levels are shown for all methylation contexts along all seven F. vesca chromosomes, using a 50 kb window. ACCT= 'Alta' crown cold-treated; ALCT= 'Alta' leaf cold-treated; FCCT= 'FDP817' crown cold-treated; FLCT= 'FDP817' leaf cold-treated; NCCT= 'NCGR1363' crown cold-treated; NLCT= 'NCGR1363' leaf cold-treated.

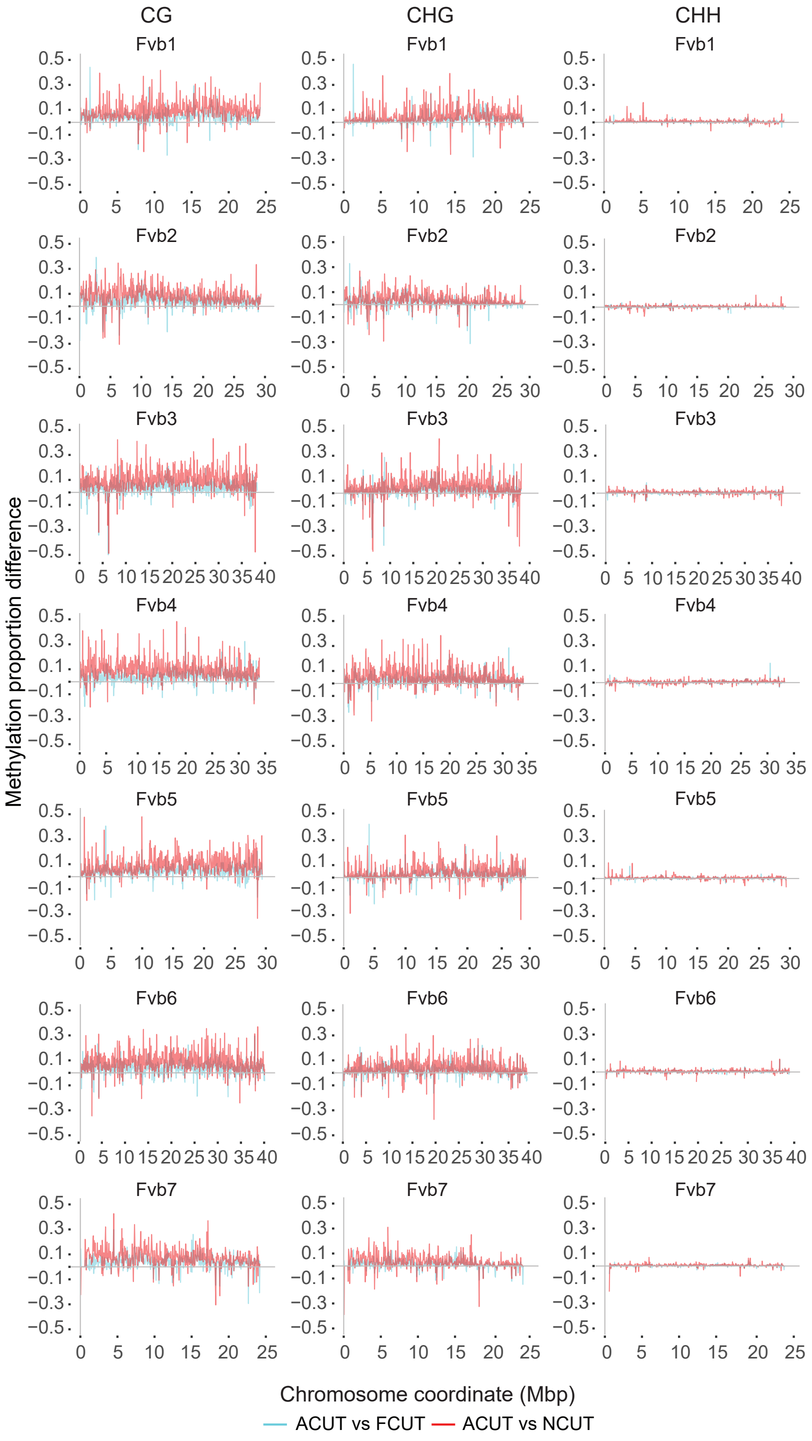


**Figure S4. DNA methylation differences in untreated crowns of ecotypes.** 'Alta' has higher basal methylation levels (methylated reads/total reads) than 'FDP817' and 'NCGR1363.' Methylation levels are shown for all methylation contexts along all seven F. vesca chromosomes, using a 50 kb window. ACUT= 'Alta' crown untreated; FCUT= 'FDP817' crown untreated; NCUT= 'NCGR1363' crown untreated.

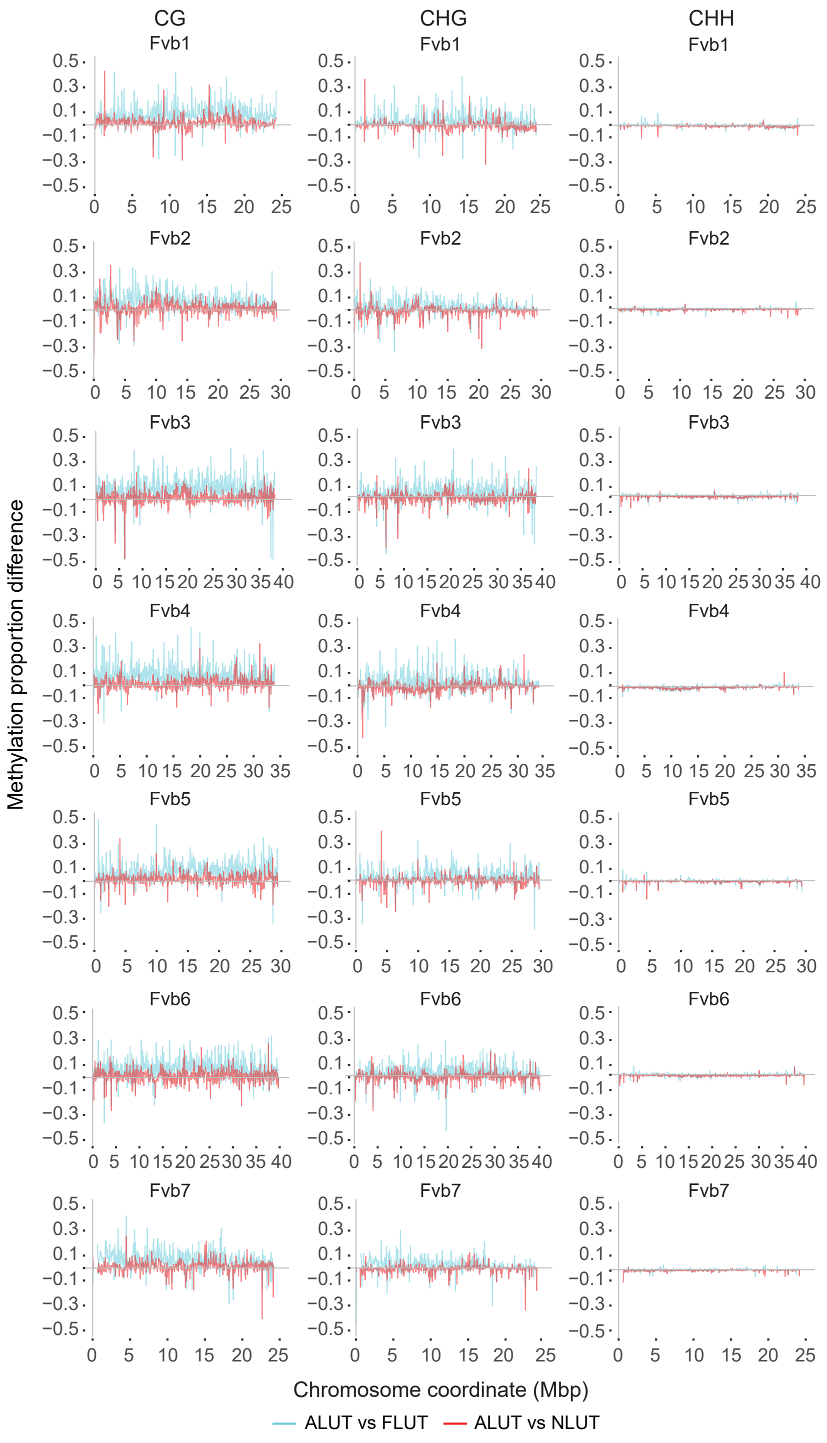


**Figure S5. DNA methylation differences in untreated leaves of ecotypes.** 'Alta' has higher basal methylation levels (methylated reads/total reads) than 'FDP817' and 'NCGR1363.' Methylation levels are shown for all methylation contexts along all seven *F. vesca* chromosomes, using a 50 kb window. ALUT= 'Alta' leaf untreated; FLUT= 'FDP817' leaf untreated; NLUT= 'NCGR1363' leaf untreated.

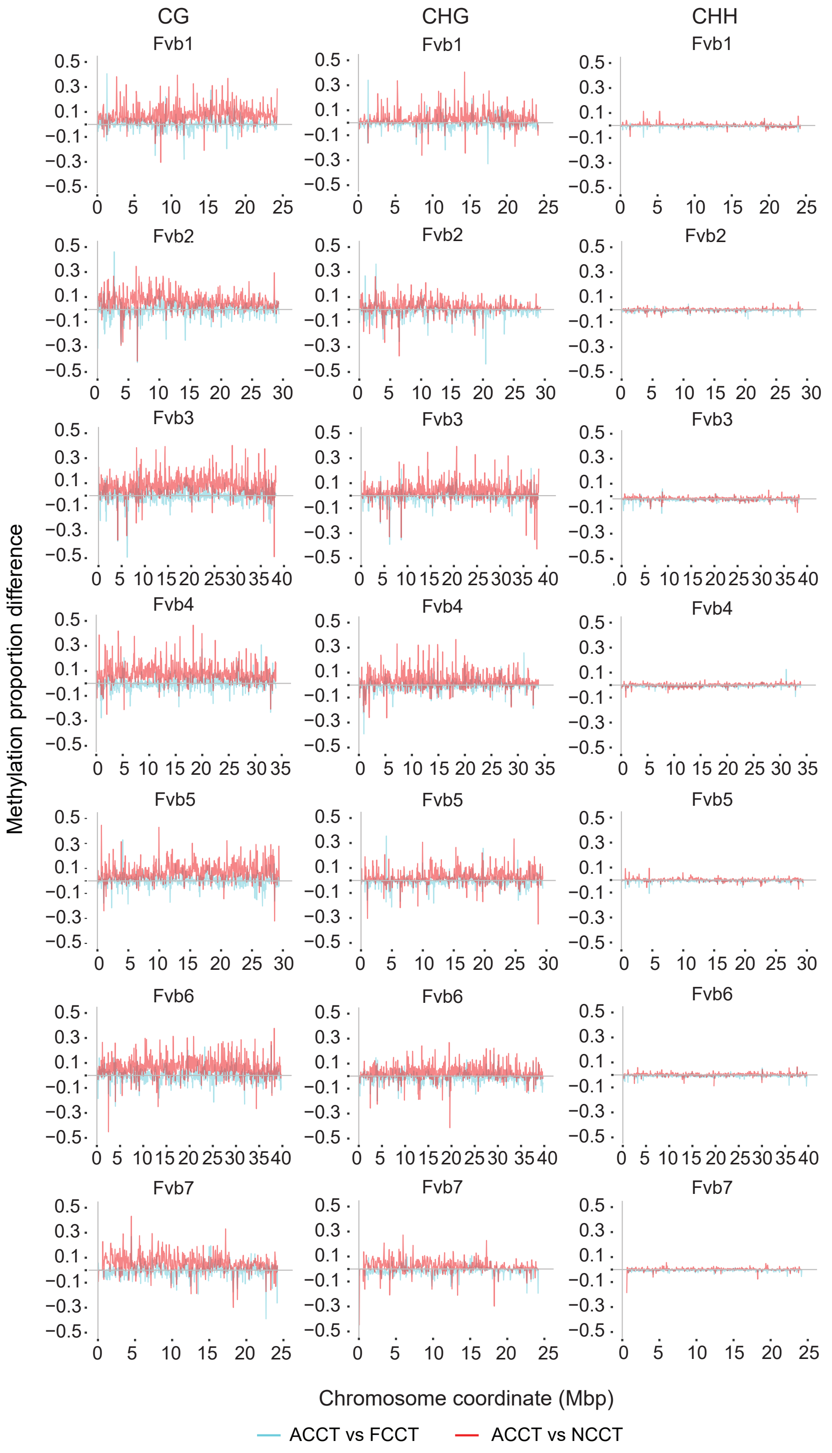


**Figure S6. DNA methylation differences in cold-treated crowns of ecotypes.** ‘Alta’ has higher methylation levels (methylated reads/total reads) than ‘FDP817’ and ‘NCGR1363.’ Methylation levels are shown for all methylation contexts along all seven *F. vesca* chromosomes, using a 50 kb window. ACCT= ‘Alta’ crown cold-treated; FCCT= ‘FDP817’ crown cold-treated; NCCT= ‘NCGR1363’ crown cold-treated.

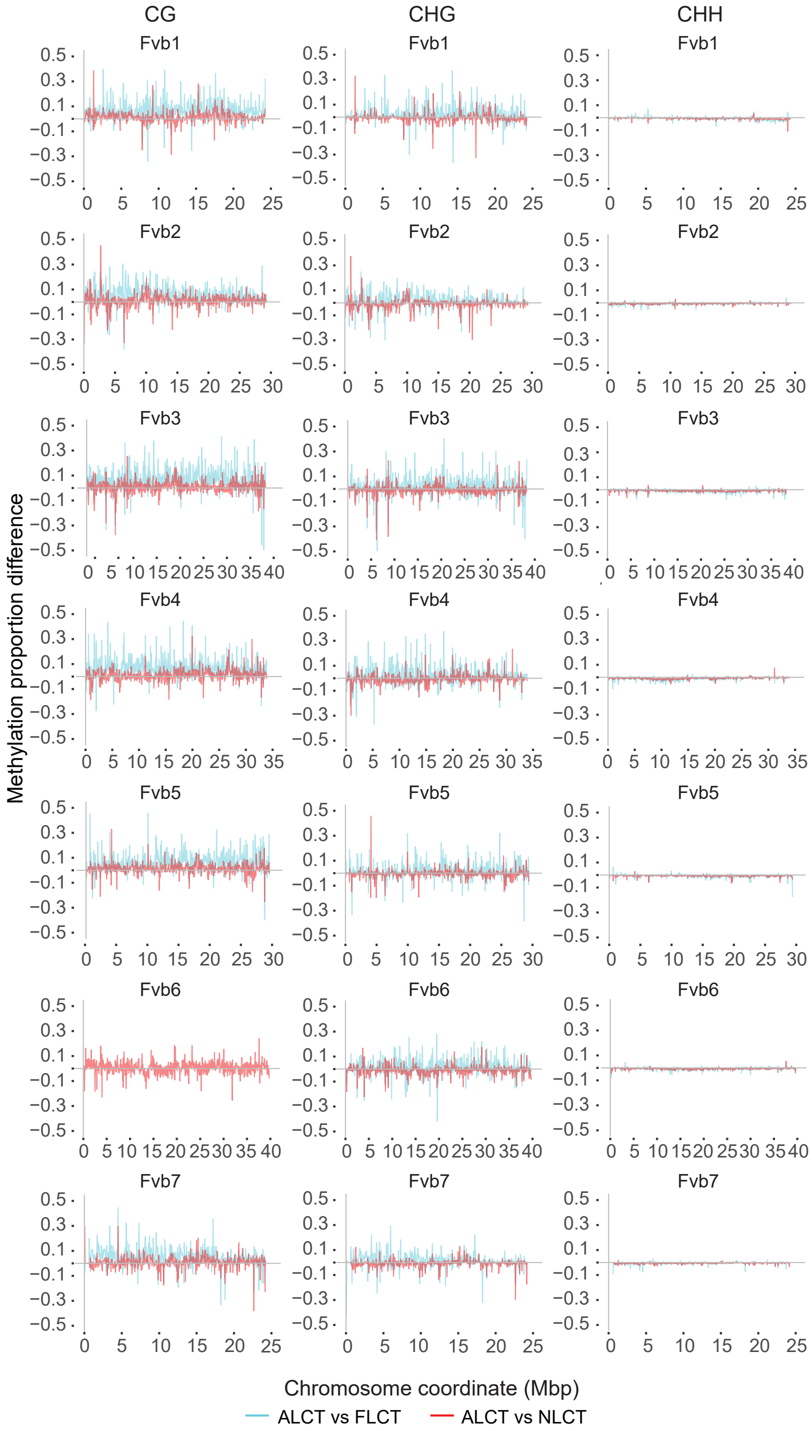


**Figure S7. DNA methylation differences in cold-treated leaves of ecotypes.** 'Alta' has higher methylation levels (methylated reads/total reads) than 'FDP817' and 'NCGR1363.' Methylation levels are shown for all methylation contexts along all seven *F. vesca* chromosomes, using a 50 kb window. ALCT= 'Alta' leaf cold-treated; FLCT= 'FDP817' leaf cold-treated; NLCT= 'NCGR1363' leaf cold-treated.

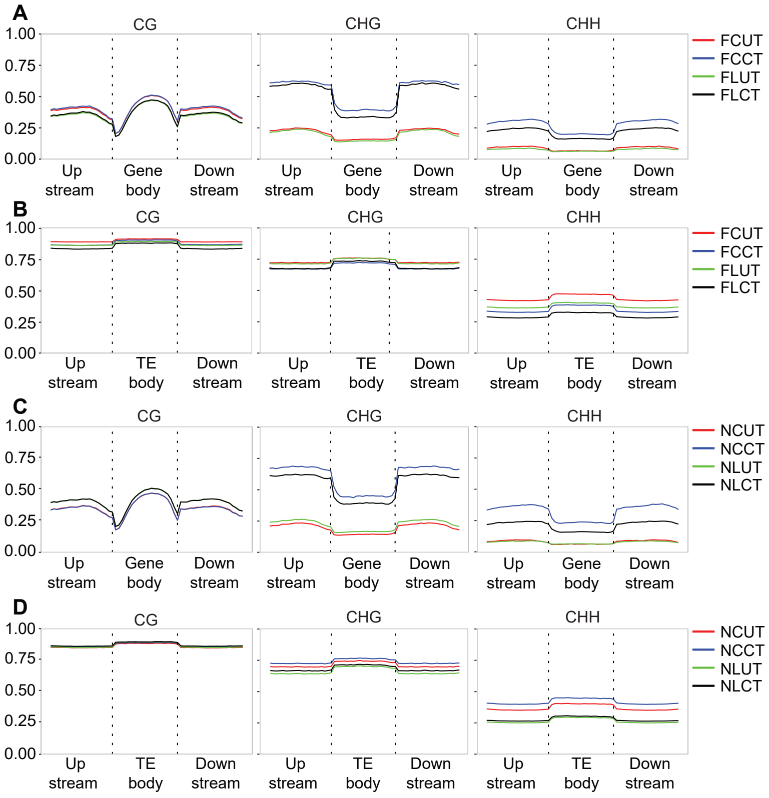


**Figure S8. DNA methylation patterns in genomic regions of 'FDP817' and 'NCGR1363' propagated at normal (UT; 20 ± 2 °C) and low (CT; 2 °C) temperature conditions for 42 days.** (A) DNA methylation patterns in CG, CHG, and CHH sites of genic regions in 'FDP817' crowns and leaves. (B) DNA methylation patterns in CG, CHG, and CHH sites of TE regions in 'FDP817' crowns and leaves. (C) DNA methylation patterns in CG, CHG, and CHH sites of genic regions in 'NCGR1363' crowns and leaves. (D) DNA methylation patterns in CG, CHG, and CHH sites of TE regions in 'NCGR1363' crowns and leaves. Plots show cold-induced methylation changes in CHG and CHH contexts in both genic regions and TE regions. TE regions have higher basal methylation levels (UT) and undergo smaller methylation changes after cold acclimation compared with genic regions. Upstream and downstream regions are 1 kb in size. Each genomic feature and regions 1 kb upstream and downstream of these were divided into 20 windows, and the average methylation level per window was plotted to illustrate global DNA methylation levels. Colored lines show different combinations of tissue and temperature: FCUT= 'FDP817' crown Untreated, FLUT= 'FDP817' leaf Untreated, FCCT= 'FDP817' crown cold-treated, FLCT= 'FDP817' leaf cold-treated, NCUT= 'NCGR1363' crown untreated, NLUT= 'NCGR1363' leaf untreated, NCCT= 'NCGR1363' crown cold-treated, NLCT= 'NCGR1363' leaf cold-treated.

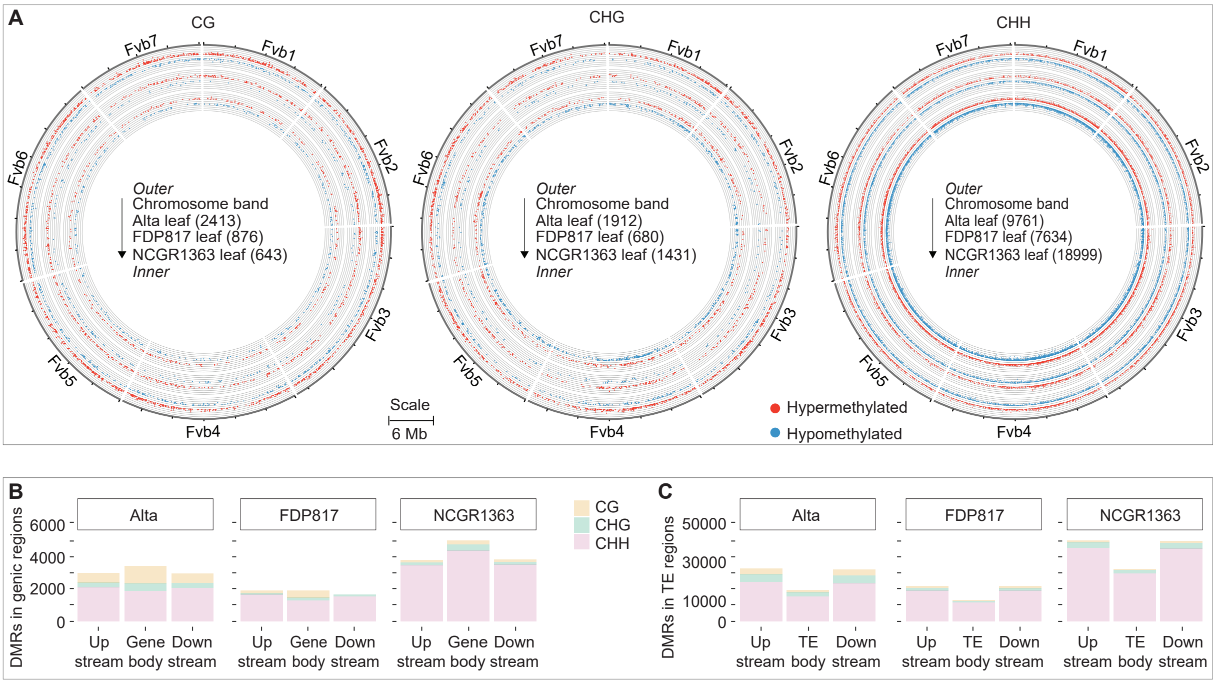


**Figure S9. DMR patterns in ecotype leaves propagated under normal (UT; 20 ± 2 °C) and low (CT; 2 °C) temperature conditions for 42 days.** (A) Distribution of differentially methylated regions at CG, CHG, and CHH contexts in all seven chromosomes of ecotypes. Numbers of DMRs in each tissue are indicated in parentheses. Red and blue dots denote hypermethylated and hypomethylated regions, respectively. (B) Number of DMRs overlapping with CG, CHG, and CHH sites in genic regions of ecotype leaves. (C) Number of DMRs overlapping with CG, CHG, and CHH sites in TE regions of ecotype leaves. 'NCGR1363' has the highest number of DMRs across genomic regions, with a majority of DMRs in the CHH context. Differential methylation in TE regions is about 10 times higher than in genic regions. Upstream and downstream regions are 1 kb in size.

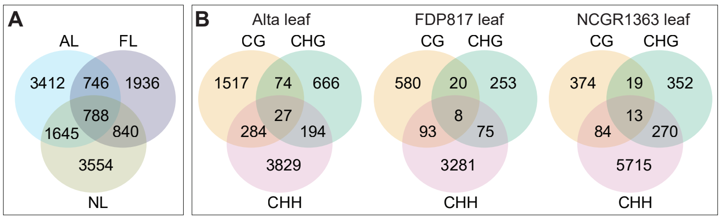


**Figure S10. Overlap of differentially methylated genes (DMGs) between 'Alta,' 'NCGR1363' and 'FDP817' leaves.** (A) Numbers of shared and unique DMGs in ecotypes: 'Alta' and 'NCGR1363' share less than half of their DMGs; most DMGs in 'FDP817' are shared; 'Alta' and 'NCGR1363' share more DMGs than either shares with 'FDP817.' (B) Numbers of genes with DMRs in either one or multiple methylation context(s). A majority of genes are methylated exclusively in one context.

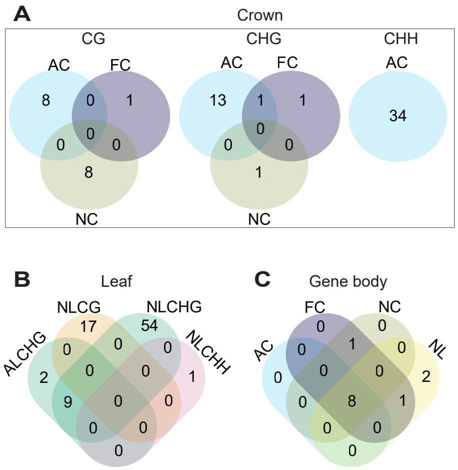


**Figure S11. Significantly enriched Gene Ontology (GO) terms among differentially methylated genes (DMGs) in different ecotypes.** Numbers of unique and shared GO terms at CG, CHG, and CHH contexts of ecotype crowns (A) and leaves (B). GO terms were mostly unique to each context and ecotype. (C) Shared and unique GO terms overlapping in gene bodies among ecotypes. Most enriched GO terms are shared by all ecotypes. AC= 'Alta' crown, AL= 'Alta' leaf, NC= 'NCGR1363' crown, NL= 'NCGR1363' leaf; FC= 'FDP817' crown.

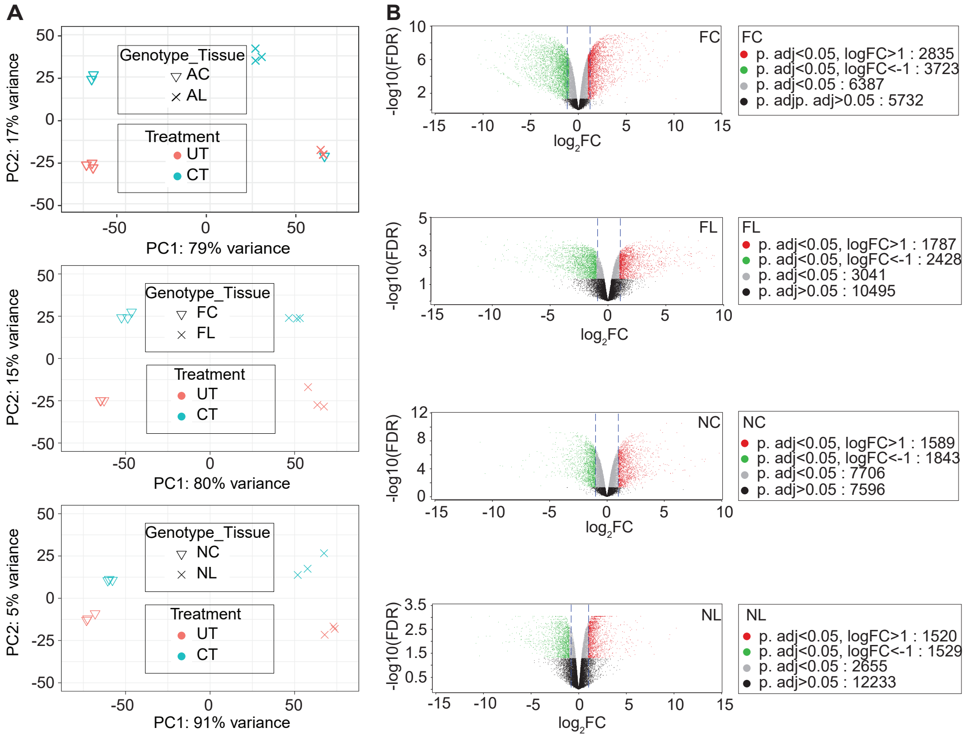


**Figure S12. Homogeneity assessment and expression profile in crowns and leaves of 'Alta', 'FDP817', and 'NCGR1363' propagated at 20 ± 2 °C (UT) and 2 °C (CT) for 42 days.** (A) Clustering of replicates into treatment groups for crowns and leaves of ecotypes propagated at control (UT; 20 ± 2 °C) and cold acclimation (CT; 2 °C) conditions for 42 days. Principal component analysis revealed homogeneity within replicates of the same treatment group, except for 'Alta' crown (CT), for which one replicate was excluded from downstream analysis due to divergence from the other two replicates. (B) Volcano plots of expressed (-log10FDR) genes versus their degree of fold change (log2FC). Significantly expressed genes (p.adjust < 0.05) are marked in green, red, and grey, with red and green denoting genes with log2FC values greater than 1 (upregulated) and less than -1 (downregulated), respectively. Black dots indicate non-significant genes with p.adjust ≥ 0.05. Points within dashed lines define -1 ≤ log2FC ≤ 1. UT= Untreated, CT= Cold-treated.

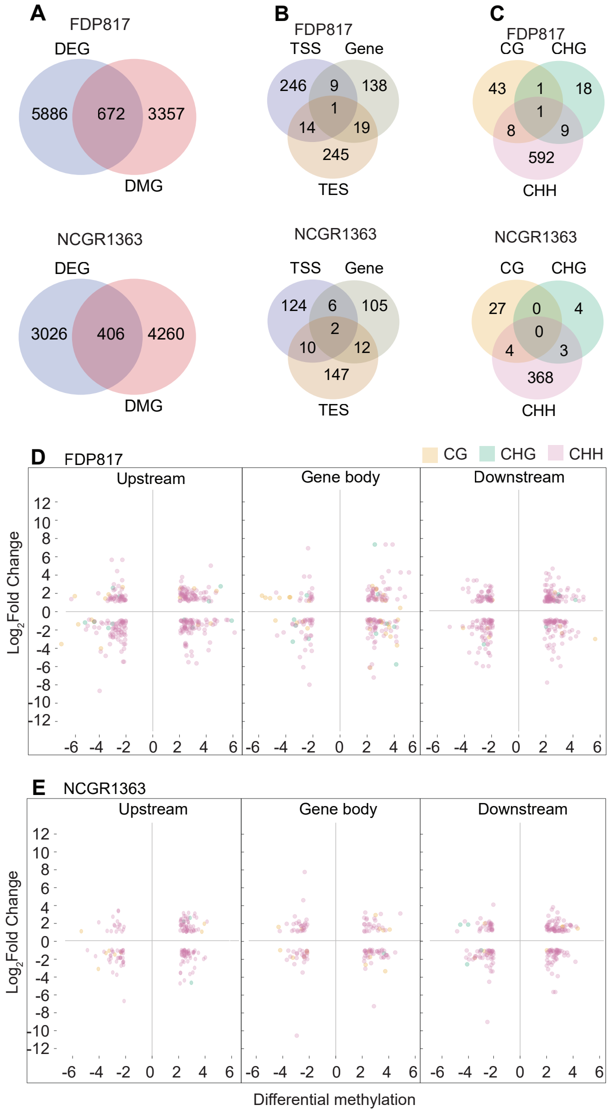


**Figure S13. Correlation of differential methylation and gene expression in crowns of ecotypes.** (A) Overlap between DEGs and DMGs in 'FDP817' and 'NCGR1363.' (B) Number of DEDMGs harboring DMRs in either one or multiple genic regions. (C) Number of DEDMGs with DMRs in one or multiple methylation contexts. A majority of DEDMGs are methylated exclusively in one context. (D, E) Differentially expressed genes associated with DMRs (DEDMRs), showing a relationship between transcript accumulation (log2fold change) and differential methylation levels (proportion difference between cold-treated and untreated samples) in CG, CHG, and CHH contexts of gene bodies and regions 1kb upstream of transcription start sites and downstream of transcription end sites in 'FDP817' and 'NCGR1363.'

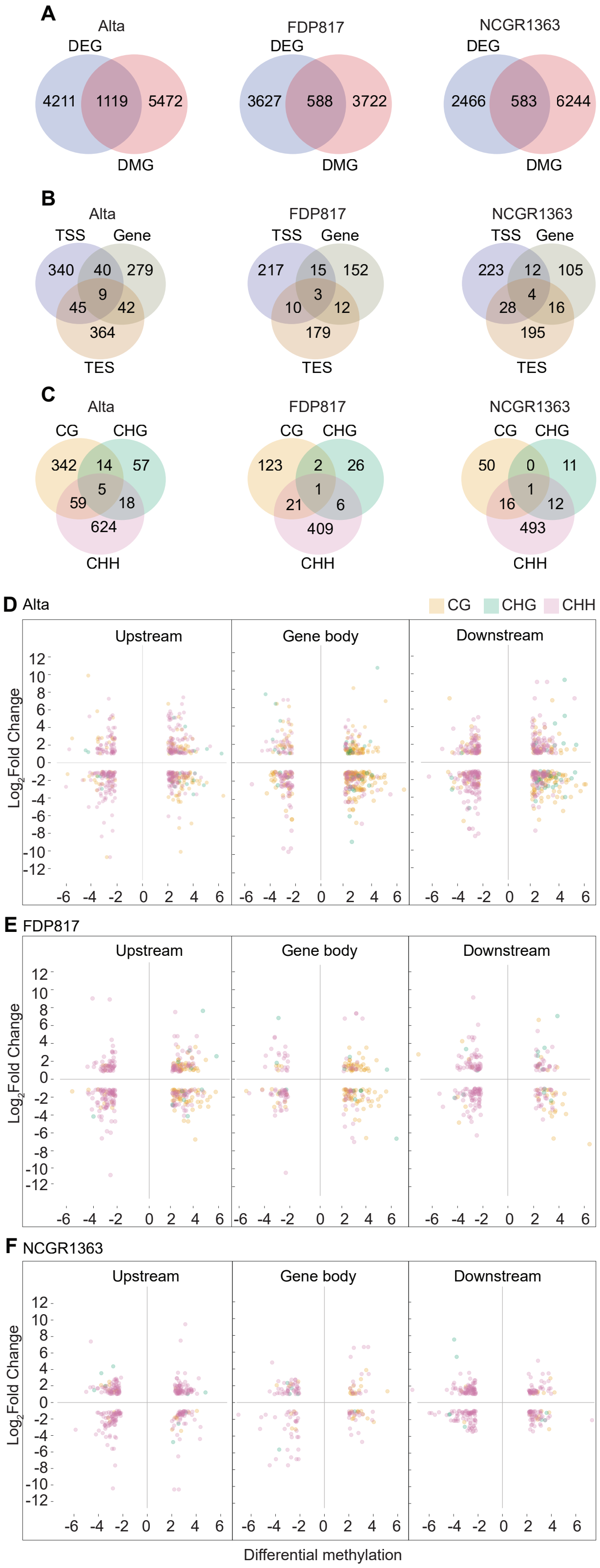


**Figure S14. Correlation of differential methylation and gene expression in leaves of ecotypes.** (A) Overlap between DEGs and DMGs in ‘Alta,’ ‘FDP817’ and ‘NCGR1363.’ (B) Number of DEDMGs harboring DMRs in either one or multiple genic regions. (C) Number of DEDMGs with DMRs in one or multiple methylation contexts. (D-F) Differentially expressed genes associated with DMRs (DEDMRs), showing relationship between transcript accumulation (log2 fold change) and DNA methylation levels (DNA methylation proportion difference between 42D cold acclimated and control samples) in CG, CHG and CHH contexts of gene bodies and regions 1kb upstream of transcription start site and downstream of transcription end site in ‘Alta,’ ‘FDP817’, and ‘NCGR1363.’

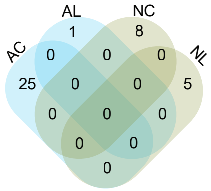


**Figure S15. Gene Ontology (GO) enrichment analysis of differentially expressed and methylated genes (DEDMGs).** GO terms enriched in DEDMGs unique to tissues among ecotypes. Most enriched GO terms are unique to ecotype and methylation context. AC= ‘Alta’ crown, NC= ‘NCGR1363’ crown, AL= ‘Alta’ leaf, NL= ‘NCGR1363’ leaf.

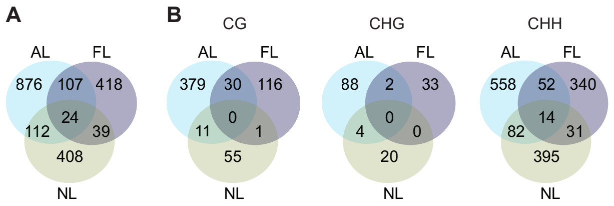


**Figure S16. Differentially expressed and differentially methylated genes (DEDMGs) in leaves of 'Alta,' 'FDP817' and 'NCGR1363.'** (A) Number of shared and unique DEDMGs in ecotypes regardless of methylation context and region. (B) Number of shared and unique DEDMGs in CG, CHG, and CHH contexts among ecotypes. Most DEDMGs are ecotype specific. AL= 'Alta' leaf, FL= 'FDP817' leaf, NL= 'NCGR1363' leaf.

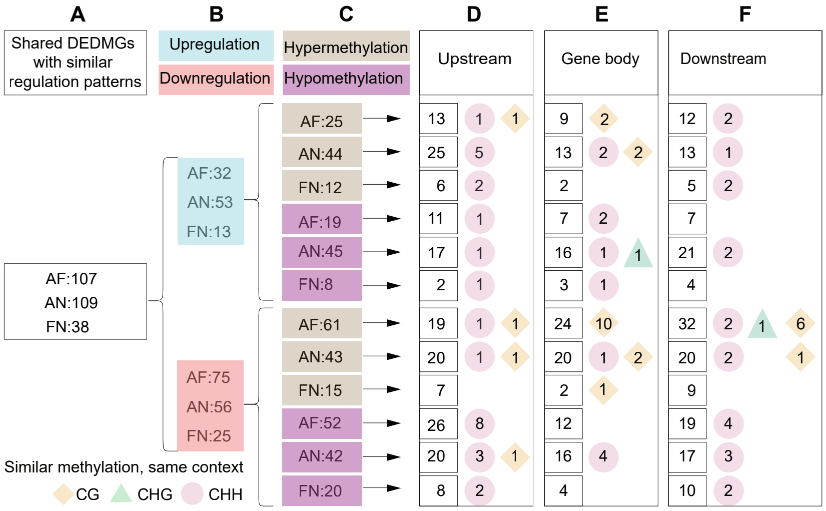


**Figure S17. Analysis of leaf DEDMGs shared between ecotypes.** (A) Numbers of shared DEDMGs differentially expressed in the same directions between ecotypes. (B) Subsets of shared DEDMGs expressed in the same directions. (C) DEDMGs from column B with differential methylation in the same directions. (D-F) Numbers of shared DEDMGs from column C, showing differential methylation in the same directions in genic regions (squares) and contexts within regions (diamond and circles) of ecotype pairs. The diamonds, triangles, and circles represent CG, CHG, and CHH methylation contexts, respectively. AF= ‘Alta’ and ‘FDP817,’ AN= ‘Alta’ and ‘NCGR1363,’ FN= ‘FDP817’ and ‘NCGR1363.’

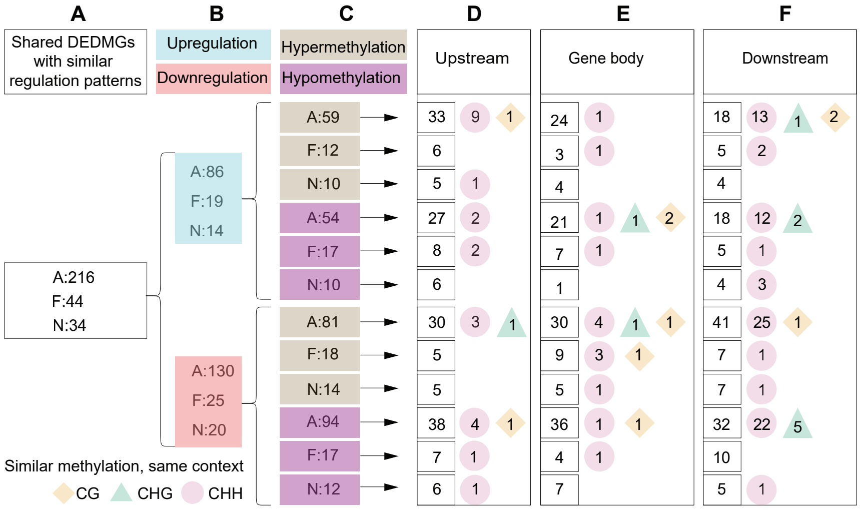


**Figure S18. Analysis of crown and leaf DEDMGs shared within ecotypes.** (A) Numbers of shared DEDMGs differentially expressed in the same directions between ecotypes. (B) Subsets of shared DEDMGs expressed in the same directions. (C) DEDMGs from column B with differential methylation in the same directions. (D-F) Numbers of shared DEDMGs from column C, showing differential methylation in the same directions in genic regions (squares) and contexts within regions (diamond and circles) of ecotype pairs. The diamonds, triangles, and circles represent CG, CHG, and CHH methylation contexts, respectively. A= 'Alta', F= 'FDP817', N= 'NCGR1363.'
