## Supplemental Tables S1-S7 for "Epigenetic plasticity is associated with enhanced tolerance to low temperature stress in woodland strawberry"

**Table S1. Genotypes of woodland strawberry used in this study.** The genotypes are collected from different regions and show different cold tolerance abilities. Estimated $LT_{50}$ values indicate freezing temperatures at which 50 percent of cold-treated plants do not survive (Davik et al., 2013). NCGR = National Clonal Germplasm Repository. CFRA 371 and CFRA1363 are the NCGR accession IDs also identified as NCGR371 and 'NCGR1363', respectively, by Davik et al. (2013). CFRA371 was also denoted as 'FDP817' by the Horticultural Research International (HRI) in the UK (Hadonou et al. 2004). USDA-ARS, United States Department of Agriculture-Agricultural Research Service. m.a.s.l= meters above sea level.

| **Ecotype** | **Estimated $LT_{50}$** | **Origin** | **Latitude (°N)** | **Longitude** | **Altitude (m.a.s.l)** | **Reference** |
|---|---|---|---|---|---|---|
| Alta | -11.6 | Alta, Norway | 69.9 | 23.0° E | 40 | Davik et al. 2013 |
| NCGR1363 | -8.8 | Bolivia | | | | Davik et al. 2013 |
| FDP817 | -7.7 | California, USA | 34.4 | 120.5° W | 15 | Davik et al. 2013 |

**Table S2. WGBS mapping statistics showing the efficiency level of mapping sequenced reads of individual ecotypes to *Fragaria vesca* version 4 (FvH4.0.a2) genome**. Mapping efficiency varied from between 51 % to 70 % at the context and treatment level across 'Alta', 'FDP817', and 'NCGR1363' ecotypes, with leaves having a higher number of mapped reads than crowns. The majority of methylated cytosines were recorded in the CG context, followed by CHG and then CHH. ACUT= 'Alta' crown untreated, ALUT= 'Alta' leaf untreated, ACCT= 'Alta' crown cold-treated, ALCT= 'Alta' leaf cold-treated, FCUT= 'FDP817' crown untreated, FLUT= 'FDP817' leaf untreated, FCCT= 'FDP817' crown cold-treated, FLCT= 'FDP817' leaf cold-treated, NCUT= 'NCGR1363' crown untreated, NLUT= 'NCGR1363' leaf untreated, NCCT= 'NCGR1363' crown cold-treated, NLCT= 'NCGR1363' leaf cold-treated.

| Sample ID | Total mapped reads (R1+R2) | Mapping efficiency (%) | Paired reads after deduplication | Number of C's analyzed | Methylated C's in CG (%) | Methylated C's in CHG (%) | Methylated C's in CHH (%) |
|---|---|---|---|---|---|---|---|
| ACUT | 143105406 | 65 | 42543361 | 1484172920 | 37.3 | 16 | 3.7 |
| ACCT | 75945752 | 52.4 | 18213058 | 644819394 | 31.6 | 13.6 | 3.1 |
| ALUT | 105541126 | 69.4 | 33712142 | 1141143031 | 38.1 | 15.7 | 2.4 |
| ALCT | 65808104 | 66.9 | 20453694 | 689451440 | 37.2 | 15.3 | 2 |
| FCUT | 46452104 | 59.6 | 12831840 | 461883873 | 30.4 | 13.3 | 3.2 |
| FCCT | 66491314 | 54.4 | 18663053 | 671081363 | 32.1 | 14.9 | 3.8 |
| FLUT | 57600452 | 62 | 16718663 | 573481828 | 30.1 | 12.4 | 2.5 |
| FLCT | 79648546 | 61.3 | 25953006 | 872940677 | 30.6 | 13 | 2.5 |
| NCUT | 82552498 | 51.5 | 19684880 | 697813893 | 26 | 11 | 2.9 |
| NCCT | 77283094 | 53 | 15937766 | 562746383 | 24.9 | 10.7 | 2.8 |
| NLUT | 85270524 | 70.2 | 27853802 | 943080822 | 37.5 | 16.1 | 3 |
| NLCT | 93853778 | 70.8 | 25725505 | 853248111 | 35.7 | 15.7 | 2.5 |

**Table S3. Number of DMRs showing methylation direction in CG, CHG, and CHH contexts of 'Alta', 'FDP817', and 'NCGR1363' tissues**. DMRs in CG and CHG contexts were called with a bin size of 100 bp, and CHH with 50 bp.

| Ecotype tissue | Context | DMR numbers | Number of hypermethylated DMRs | Number of hypomethylated DMRs | Total DMRs | Total DMRs |
|---|---|---|---|---|---|---|
| Alta crown | CG | 2149 | 791 | 1358 | 23193 | 37279 |
| | CHG | 2707 | 459 | 2248 | | |
| | CHH | 18337 | 9998 | 8339 | | |
| Alta leaf | CG | 2413 | 1668 | 745 | 14086 | |
| | CHG | 1912 | 1393 | 519 | | |
| | CHH | 9761 | 4086 | 5675 | | |
| FDP817 crown | CG | 281 | 162 | 119 | 10636 | 19826 |
| | CHG | 646 | 419 | 227 | | |
| | CHH | 9709 | 5898 | 3811 | | |
| FDP817 leaf | CG | 876 | 660 | 216 | 9190 | |
| | CHG | 680 | 466 | 214 | | |
| | CHH | 7634 | 3148 | 4486 | | |
| NCGR1363 crown | CG | 426 | 204 | 222 | 11473 | 32546 |
| | CHG | 311 | 145 | 166 | | |
| | CHH | 10736 | 5991 | 4745 | | |
| NCGR1363 leaf | CG | 643 | 337 | 306 | 21073 | |
| | CHG | 1431 | 535 | 896 | | |
| | CHH | 18999 | 5592 | 13407 | | |

**Table S4. Number of DMRs in genic regions and contexts 'Alta', 'FDP817', and 'NCGR1363' tissues.**

| Tissue | Region | Context | Alta | FDP817 | NCGR1363 | Total DMRs | | |
|---|---|---|---|---|---|---|---|---|
| | | | | | | Alta | FDP817 | NCGR1363 |
| Crown | Upstream | CG | 492 | 65 | 87 | 4463 | 1819 | 2175 |
| | | CHG | 379 | 96 | 50 | | | |
| | | CHH | 3592 | 1658 | 2038 | | | |
| | Gene body | CG | 1060 | 139 | 187 | 4745 | 2300 | 1923 |
| | | CHG | 736 | 208 | 83 | | | |
| | | CHH | 2949 | 1953 | 1653 | | | |
| | Downstream | CG | 536 | 64 | 70 | 4570 | 1771 | 2200 |
| | | CHG | 382 | 71 | 57 | | | |
| | | CHH | 3652 | 1636 | 2073 | | | |
| Leaf | Upstream | CG | 610 | 158 | 152 | 3040 | 1940 | 3871 |
| | | CHG | 284 | 106 | 181 | | | |
| | | CHH | 2146 | 1676 | 3538 | | | |
| | Gene body | CG | 1077 | 456 | 240 | 3493 | 1946 | 5078 |
| | | CHG | 503 | 182 | 398 | | | |
| | | CHH | 1913 | 1308 | 4440 | | | |
| | Downstream | CG | 592 | 168 | 129 | 3008 | 1840 | 3880 |
| | | CHG | 309 | 102 | 188 | | | |
| | | CHH | 2107 | 1570 | 3563 | | | |

**Table S5. Number of DMRs in TE regions and contexts 'Alta', 'FDP817', and 'NCGR1363' tissues.**

| Tissue | Region | Context | Alta | FDP817 | NCGR1363 | Total DMRs | | |
|---|---|---|---|---|---|---|---|---|
| | | | | | | Alta | FDP817 | NCGR1363 |
| Crown | Upstream | CG | 2672 | 394 | 592 | 48571 | 21445 | 24571 |
| | | CHG | 5742 | 1267 | 587 | | | |
| | | CHH | 40157 | 19784 | 23392 | | | |
| | TE body | CG | 1137 | 170 | 262 | 28475 | 13056 | 14178 |
| | | CHG | 3130 | 757 | 328 | | | |
| | | CHH | 24208 | 12129 | 13588 | | | |
| | Downstream | CG | 2667 | 332 | 591 | 48502 | 21518 | 24223 |
| | | CHG | 5734 | 1300 | 628 | | | |
| | | CHH | 40101 | 19886 | 23004 | | | |
| Leaf | Upstream | CG | 3140 | 1047 | 927 | 27898 | 18576 | 42488 |
| | | CHG | 4095 | 1434 | 3134 | | | |
| | | CHH | 20663 | 16095 | 38427 | | | |
| | TE body | CG | 1288 | 473 | 459 | 16549 | 11117 | 27378 |
| | | CHG | 2164 | 731 | 1757 | | | |
| | | CHH | 13097 | 9913 | 25162 | | | |
| | Downstream | CG | 3004 | 1057 | 982 | 27139 | 18527 | 42205 |
| | | CHG | 4171 | 1392 | 3057 | | | |
| | | CHH | 19964 | 16078 | 38166 | | | |

**Table S6. RNA-seq and mapping statistics of reads generated from Illumina sequencing of leaves and crowns of untreated (UT) and cold-treated (CT) 'Alta', 'FDP817', and 'NCGR1363.'**

| Ecotype/Treatment | Biological replicate | Raw reads | | | | Total mapped reads | Mapping efficiency (%) |
|---|---|---|---|---|---|---|---|
| | | Technical replicate 1 | | Technical replicate 2 | | | |
| | | Read 1 | Read 2 | Read 1 | Read 2 | (Read1+Read2) | |
| Alta crown/UT | 1 | 10,478,081 | 10,478,081 | 10,637,517 | 10,637,517 | 15,158,104 | 72.03 |
| | 2 | 9,910,590 | 9,910,590 | 10,063,588 | 10,063,588 | 14,276,169 | 71.72 |
| | 3 | 9,858,159 | 9,858,159 | 9,999,944 | 9,999,944 | 14,071,678 | 71.23 |
| Alta crown/CT | 1 | 9,480,672 | 9,480,672 | 9,606,454 | 9,606,454 | 14,017,573 | 73.88 |
| | 2 | 10,842,665 | 10,842,665 | 11,003,499 | 11,003,499 | 16,681,370 | 76.7 |
| | 3 | 9,135,103 | 9,135,103 | 9,245,308 | 9,245,308 | 13,503,240 | 73.85 |
| Alta leaf/UT | 1 | 10,993,941 | 10,993,941 | 11,157,680 | 11,157,680 | 16,886,737 | 76.6 |
| | 2 | 8,766,695 | 8,766,695 | 8,919,539 | 8,919,539 | 12,891,082 | 73.26 |
| | 3 | 9,664,239 | 9,664,239 | 9,802,915 | 9,802,915 | 13,969,748 | 72.53 |
| Alta leaf/CT | 1 | 10,087,424 | 10,087,424 | 10,233,997 | 10,233,997 | 14,010,919 | 69.21 |
| | 2 | 8,555,672 | 8,555,672 | 8,659,907 | 8,659,907 | 12,820,894 | 74.88 |
| | 3 | 9,775,313 | 9,775,313 | 9,728,615 | 9,728,615 | 14,351,545 | 73.92 |
| NCGR1363 crown/UT | 1 | 9,889,704 | 9,889,704 | 10,022,153 | 10,022,153 | 15,971,080 | 80.56 |
| | 2 | 9,139,541 | 9,139,541 | 9,267,654 | 9,267,654 | 14,978,207 | 81.74 |
| | 3 | 8,828,759 | 8,828,759 | 8,947,664 | 8,947,664 | 14,387,403 | 81.25 |
| NCGR1363 crown/CT | 1 | 10,612,821 | 10,612,821 | 10,762,700 | 10,762,700 | 16,846,139 | 79.06 |
| | 2 | 8,759,106 | 8,759,106 | 8,882,593 | 8,882,593 | 13,793,332 | 78.45 |
| | 3 | 11,161,853 | 11,161,853 | 11,311,207 | 11,311,207 | 17,312,074 | 77.27 |
| NCGR1363 leaf/UT | 1 | 12,508,529 | 12,508,529 | 12,718,280 | 12,718,280 | 18,490,791 | 73.68 |
| | 2 | 10,356,125 | 10,356,125 | 10,473,971 | 10,473,971 | 15,973,289 | 77.08 |
| | 3 | 9,571,707 | 9,571,707 | 9,709,069 | 9,709,069 | 14,716,125 | 76.62 |
| NCGR1363 leaf/CT | 1 | 9,344,439 | 9,344,439 | 9,497,993 | 9,497,993 | 14,023,381 | 74.78 |
| | 2 | 9,301,643 | 9,301,643 | 9,415,372 | 9,415,372 | 14,799,572 | 79.32 |
| | 3 | 12,296,804 | 12,296,804 | 12,517,522 | 12,517,522 | 18,249,248 | 73.94 |
| FDP817 crown/UT | 1 | 10,547,781 | 10,547,781 | 10,716,835 | 10,716,835 | 16,739,203 | 79.06 |
| | 2 | 9,948,115 | 9,948,115 | 10,090,905 | 10,090,905 | 11,953,981 | 75.89 |
| | 3 | 10,996,670 | 10,996,670 | 11,147,083 | 11,147,083 | 17,185,448 | 77.94 |
| FDP817 crown/CT | 1 | 8,626,593 | 8,626,593 | 8,746,058 | 8,746,058 | 13,066,249 | 75.52 |
| | 2 | 7,850,356 | 7,850,356 | 7,973,287 | 7,973,287 | 11,953,981 | 75.89 |

| | | | | | | | |
|---|---|---|---|---|---|---|---|
| | 3 | 6,818,802 | 6,818,802 | 6,894,401 | 6,894,401 | 10,427,564 | 76.51 |
| FDP817 leaf/UT | 1 | 8,655,927 | 8,655,927 | 8,748,719 | 8,748,719 | 12,912,282 | 74.57 |
| | 2 | 9,192,473 | 9,192,473 | 9,332,892 | 9,332,892 | 14,055,222 | 76.26 |
| | 3 | 9,592,618 | 9,592,618 | 9,687,395 | 9,687,395 | 14,366,833 | 74.76 |
| FDP817 leaf/CT | 1 | 9,167,702 | 9,167,702 | 9,294,709 | 9,294,709 | 13,141,112 | 71.45 |
| | 2 | 10,349,989 | 10,349,989 | 10,501,761 | 10,501,761 | 15,097,226 | 72.75 |
| | 3 | 9,314,075 | 9,314,075 | 9,418,073 | 9,418,073 | 12,784,000 | 68.49 |

**Table S7. Number of DEDMGs in regions and contexts of ecotypes and probability of overlap between DEGs and DMGs in tissues.** (A) Total number of DMGs, DEGs, and DEDMGs. (B) Numbers of DEDMGs in genic regions and contexts. (C) Probability of overlap between DEGs and DMGs in ecotypes. Representation factor is the number of overlapping genes divided by the expected number of overlapping genes drawn from two independent groups. A representation factor > 1 indicates more overlap than expected of two independent groups, a representation factor < 1 indicates less overlap than expected, and a representation factor of 1 indicates that the two groups overlap by the number of genes expected for independent groups of genes.

**A**

| Ecotype | Tissue | Number of DMGs | Number of DEGs | DEDMGs | % DEDMGs |
|---|---|---|---|---|---|
| Alta | crown | 8891 | 6098 | 1494 | 16.80 |
| | leaf | 6591 | 5330 | 1119 | 16.98 |
| FDP817 | crown | 4029 | 6558 | 672 | 16.68 |
| | leaf | 4310 | 4215 | 588 | 13.64 |
| NCGR1363 | crown | 4666 | 3432 | 406 | 8.70 |
| | leaf | 6827 | 3049 | 583 | 8.54 |

**B**

| Ecotype | Tissue | Context | Upstream | Gene body | Downstream | Total DEDMGs | | |
|---|---|---|---|---|---|---|---|---|
| | | | | | | Upstream | Gene body | Downstream |
| Alta | Crown | CG | 78 | 228 | 129 | 640 | 497 | 705 |
| | | CHG | 44 | 58 | 54 | | | |
| | | CHH | 518 | 211 | 522 | | | |
| | Leaf | CG | 118 | 206 | 127 | 449 | 390 | 479 |
| | | CHG | 22 | 34 | 41 | | | |
| | | CHH | 309 | 150 | 311 | | | |
| FDP817 | Crown | CG | 14 | 32 | 9 | 275 | 173 | 280 |
| | | CHG | 11 | 11 | 8 | | | |
| | | CHH | 250 | 130 | 263 | | | |
| | Leaf | CG | 29 | 88 | 34 | 251 | 189 | 208 |
| | | CHG | 13 | 11 | 13 | | | |
| | | CHH | 209 | 90 | 161 | | | |
| NCGR1363 | Crown | CG | 8 | 15 | 8 | 143 | 127 | 175 |
| | | CHG | 3 | 1 | 3 | | | |
| | | CHH | 132 | 111 | 164 | | | |
| | Leaf | CG | 20 | 31 | 17 | 275 | 139 | 247 |

| | | | | | |
|---|---|---|---|---|---|
| | | CHG | 9 | 6 | 9 |
| | | CHH | 246 | 102 | 221 |

**C**

| Ecotype | Tissue | p-value | Representation factor |
|---|---|---|---|
| Alta | Crown | 5.173e-04 | 0.9 |
| Alta | Leaf | 7.760e-04 | 1.1 |
| FDP817 | Crown | 2.100e-06 | 0.9 |
| FDP817 | Leaf | 0.005 | 1.1 |
| NCGR1363 | Crown | 2.472e-04 | 0.9 |
| NCGR1363 | Leaf | 0.083 | 1.0 |
